# Engineering a promiscuous pyrrolysyl-tRNA synthetase by a high throughput FACS screen

**DOI:** 10.1101/229054

**Authors:** Adrian Hohl, Ram Karan, Anstassja Akal, Dominik Renn, Xuechao Liu, Alaguraj Dharamarajnadar, Seema Ghoprade, Michael Groll, Magnus Rueping, Jörg Eppinger

## Abstract

The Pyrrolysyl-tRNA synthetase (PylRS) and its cognate tRNA^Pyl^ are used to facilitate the incorporation of non-canonical amino acids (ncAAs) into the genetic code of bacterial and eukaryotic cells by orthogonally reassigning the amber codon. Currently, the incorporation of new ncAAs requires a cumbersome engineering process composed of several positive and negative selection rounds to select the appropriate PylRS/tRNA^Pyl^ pair. Our fast and sensitive engineering approach required only a single FACS selection round to identify 110 orthogonal PylRS variants for the aminoacylation of 20 ncAAs. Pocket-substrate relationship from these variants led to the design of a highly promiscuous PylRS (HpRS), which catalyzed the aminoacylation of 31 structurally diverse lysine derivatives bearing clickable, fluorinated, fluorescent, and biotinylated entities. The high speed and sensitivity of our approach provides a competitive alternative to existing screening methodologies, and delivers insights into the complex PylRS-substrate interactions to facilitate the generation of additional promiscuous variants.

## Introduction

Site-specific incorporation of non-canonical amino acids (ncAAs) is a powerful tool to implement novel functions into the proteome, providing a plethora of opportunities to probe and engineer the protein structure (1–5). The central element of an effective and selective ncAA incorporation system is a heterogeneous aminoacyl-tRNA synthetase/tRNA (aaRS/tRNA) pair that does not cross-react with the original aaRSs, tRNAs, or canonical amino acids of the host cell (6). The pyrrolysyl-tRNA synthetase/tRNA^Pyl^ pair (PylRS/tRNA^Pyl^) has become a popular choice for the genetic code expansion (7, 8). This success is mainly attributed to the substrate flexibility of the active site (9, 10), the orthogonality in prokaryotic and eukaryotic cells (11), and the inherent suppression of the amber codon (8). Although over 150 ncAAs were genetically encoded during the past decade, these structures were mainly based on a few core motifs (1). Several structural functionalities still remain elusive, e.g., glycosylated or biotinylated ncAAs. The main factor limiting further expansion of the genetic code is the complex and slow process of engineering suitable aaRS/tRNA pairs. The engineering process requires the construction of large mutant libraries from which all non-functional and non-orthogonal aaRS variants must be eliminated through alternating rounds of positive and negative selection (2).

In most cases, positive and negative selection screens based on antibiotic resistance or toxic gene expression are carried out as dead-and-alive assays on agar plates (2, 12, 13). However, in this approach, the selection conditions are hardly tunable over a broad dynamic range (5, 6). Therefore, a beneficial aaRS might be accidentally eliminated due to slow cell growth and insufficient expression of the antibiotic resistance (14). The aaRS mediated ncAA incorporation in a fluorescent reporter protein provides a more sensitive and quantitative read-out (15).

Screens based on fluorescence-activated cell sorting (FACS) have been successfully applied to evolve the aaRS specificity (16–18). However, the reported experimental setups were not optimized for the intrinsically orthogonal PylRS and involved several negative and positive selection rounds, resulting in a cumbersome FACS screening process. For example, seven alternating rounds of positive and negative selection screens were required to sort tyrosyl-tRNA synthetases (TyrRS) with specificity for O-methyl-L-tyrosine (17). Since TyrRS naturally aminoacylates the canonical amino acid tyrosine, the negative selection screen was necessary to exclude the non-orthogonal variants.

Here, we describe a FACS-based screen that takes advantage of the orthogonality of the PylRS/tRNA^Pyl^ pair and involves only a single, positive selection to identify functional PylRS variants. Unlike previously reported screens that sought aaRS/tRNA pairs with high activity for a specific ncAA, our approach identifies a substantial number of PylRS variants for the same ncAA. We then used the information on pocket-substrate relationship from these variants to engineer a PylRS that aminoacylates a broad range of lysine derivatives. We term this variant “highly promiscuous pyrrolysyl-tRNA synthetase” (HpRS).

## Results

Directed evolution of proteins has emerged as the method of choice to improve or alter properties of enzymes (19). Usually, three elements determine the efficiency of the screening process: i) library design, ii) choice of reporter and iii) screening methodology. All three should be compatible as well as to be feasible in the same order of magnitude to avoid becoming a bottleneck.

### Library design

We based the design of our mutant library on a homology model of the *Methanosarcina barkeri (M. barkeri)* PylRS (Figure 1A). Since the mutation Y349F is known to enhance the aminoacylation efficiency of PylRS (9), this mutation was included in the mutant library by default. The N311 position plays an important role as a selection filter in the substrate recognition of *M. barkeri* PylRS. While asparagine forms a H-bond with the carbamate group of ncAAs and facilitates their aminoacylation, the 20 canonical amino acids are excluded (20, 21). By retaining this position, we maintained the orthogonality of the active site; and thus, were able to eliminate the negative selection in our screen. The mutation V370R was previously shown to allow the aminoacylation of norbornene amino acids by PylRS, due to a small shift of the *β*7-*β*8 hairpin caused by an H-bond between arginine and the backbone carbonyl of F349 and D351 (22). In line with these observations, the variability of the library at position 370 was limited to valine and the positively charged H-bond donors, arginine and lysine. The selected positions 271, 274, 313, 315, 378 cover most of the binding pocket’s surface and are in direct contact with the substrate. Therefore, the introduction of smaller residues at these positions possibly increased the volume of the pocket and allowed the aminoacylation of larger ncAAs.

**Figure 1.**
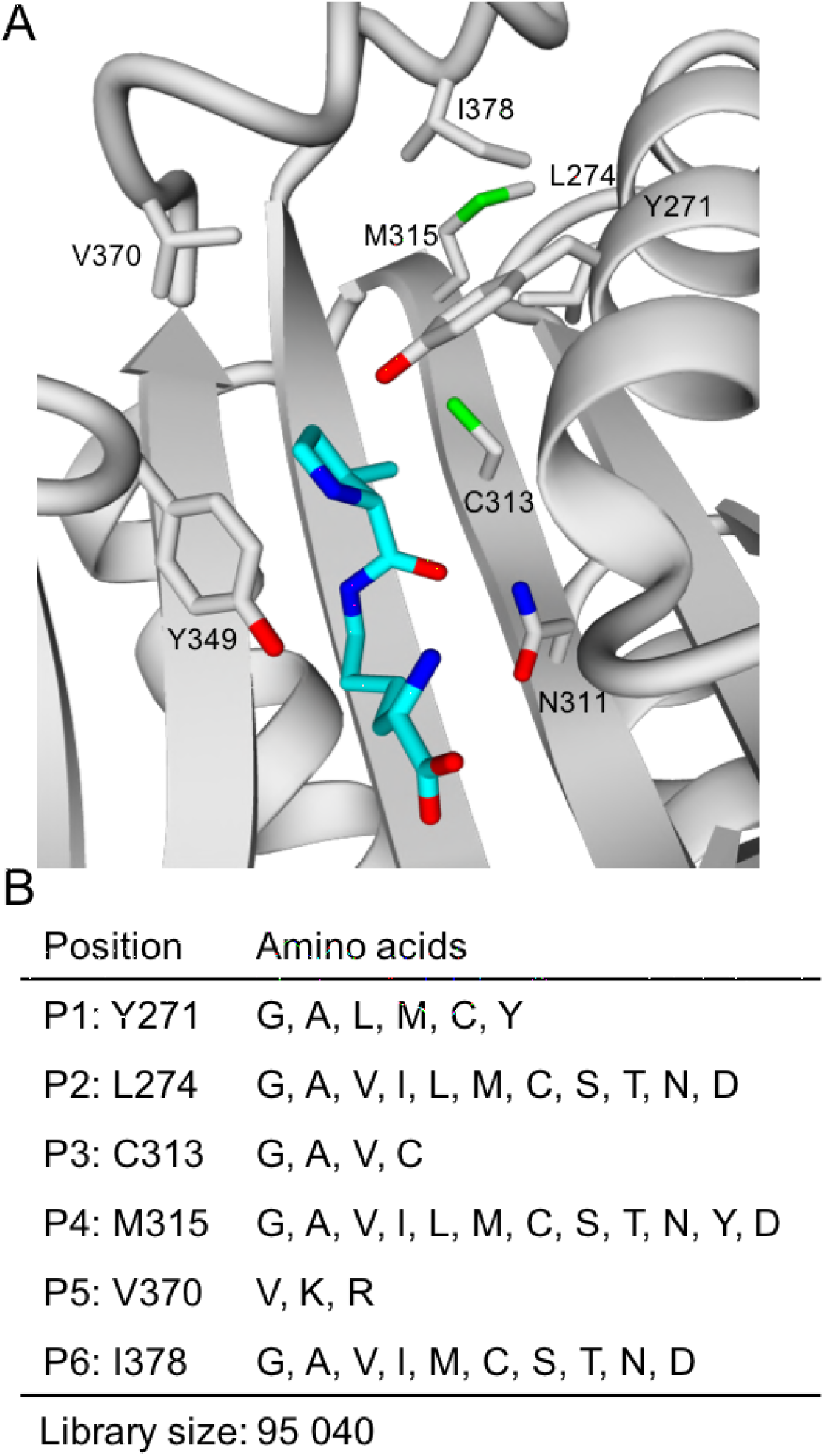
Design of *Methanosarcina barkeri* PylRS mutant library. (A) Homology model of PylRS in complex with pyrrolysine (cyan) in the binding pocket. (B) Mutation scheme of the performed mutant library. The mutation Y349F was included in the mutant library by default.

Based on these structural considerations, we selected the six amino acids Y271, L274, C313, M315, V370, and I378 to create a relatively small, focused PylRS library with 9.5x10^4^ members (Figure 1B). The library was non-codon redundant and the codon usage was optimized for *E. coli* to ensure the equal representation and expression level of all library variants. Correspondingly, 4.4x10^5^ mutants had to be screened to cover 99% of all possible sequences within the mutant library (23).

### Choice of reporter

Fluorescence-based screens provide a faster and more sensitive detection of ncAA incorporation than cell assays on selection media (15, 17). In our experiment, we chose the bright, cyan monomeric teal fluorescent protein 1 (mTFP1) (24) to report the ncAA incorporation. mTFP1 bears an amber codon at the permissive position 128. The aminoacylation of an ncAA by a corresponding PylRS/tRNA^Pyl^ pair results in the suppression of the amber codon and a bright fluorescent signal upon excitation at 462 nm from the maturated mTFP1. In contrast, the lack of a charged tRNA^Pyl^ leads to a truncated, non-fluorescent mTFP1. Thus, the aminoacylation efficiency can be quantified from fluorescence measurements (Figure 2A).

**Figure 2.**
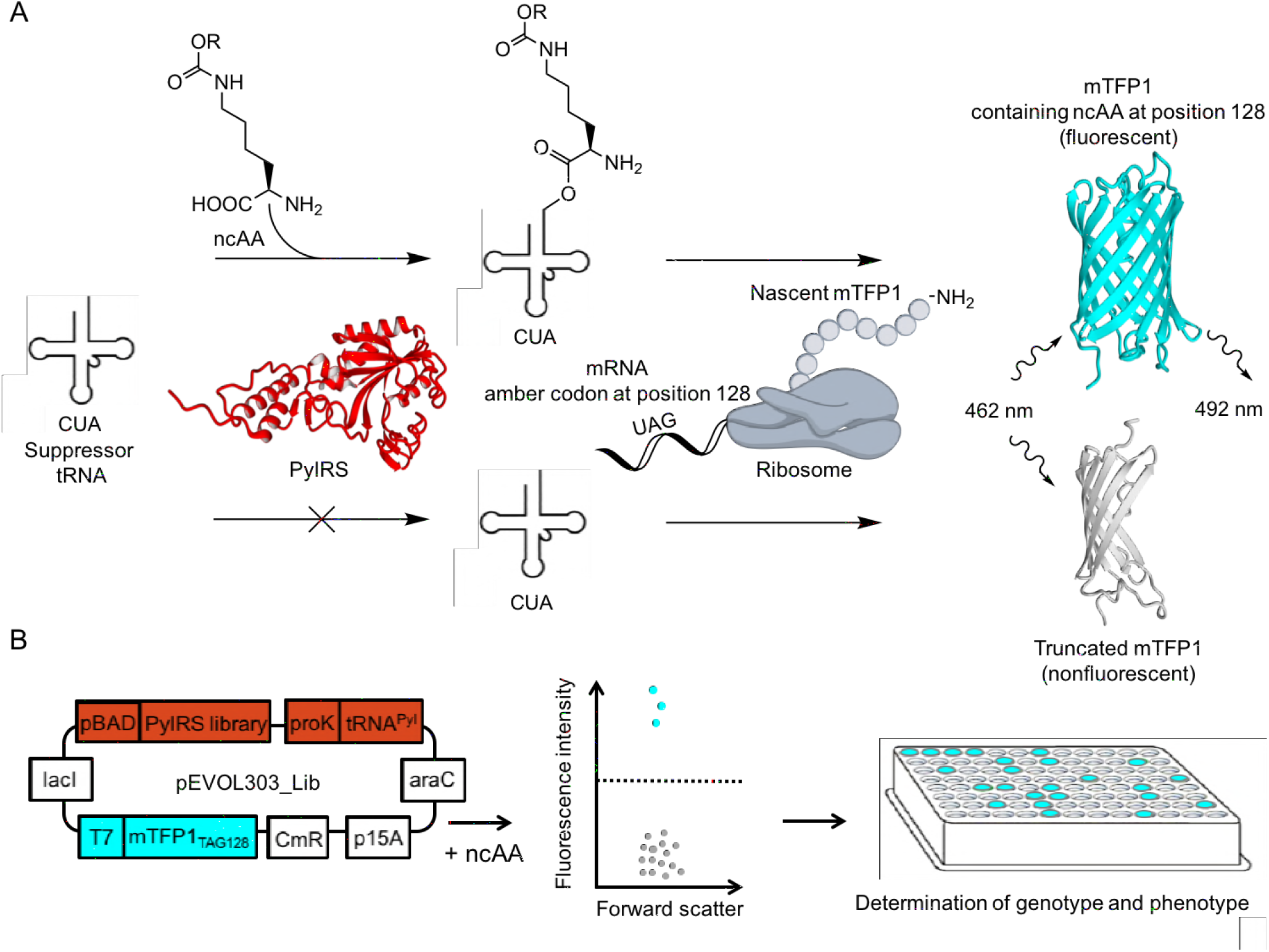
FACS-based screen of ncAA incorporation. (A) Suppression of the amber codon at position 128 in mTFP1 by a charged tRNA^Pyl^ results in a fluorescent product (ex. 462 nm, em. 492 nm). Rejection of an ncAA by PylRS leads to a truncated, nonfluorescent product. (B) Cells transformed with the plasmid pEVOL303_Lib are screened for ncAA incorporation by mTFP1-based fluorescence intensity and forward scatter using FACS. Fluorescent cells (cyan) exceeding a set threshold are sorted separately from nonfluorescent cells (grey) for sequencing on a 96-well plate. pEVOL303_Lib encodes for the PylRS library, tRNA^Pyl^ and mTFP1_TAG128_. p15A origin of replication (p15A), *araC* repressor gene (araC), chloramphenicol acetyltransferase marker (CmR), *lac* repressor (lacI), T7 promoter (T7), proK promoter (proK), araBAD promoter (pBAD).

A one-plasmid system, pEVOL303_Lib (Figure 2B), combined the required genetic components for the fluorescent screen – mTFP1_TAG128_, tRNA^Pyl^, and the PylRS library. The pEVOL303_Lib harbored the independently inducible promoter systems pBAD and T7 downstream of the PylRS and mTFP1_TAG128_.

### Fluorescence-activated cell sorting screening

*E. coli* cells containing pEV0L303_Lib were grown both in the absence and presence of ncAAs **1–23** for 16 h (Figure 3). Then, the forward scatter and fluorescent signals of each cell were analyzed by Fluorescence-activated cell sorting (FACS). The fluorescent signals of the *E. coli* cells grown without ncAAs were negligible, i.e., the PylRS library displayed no reactivity with the canonical amino acids (Figure 4A, left plot). In the presence of ncAA **1**, on the other hand, several cells depicted significant fluorescent signals (Figure 4A, right plot). Notably, the fluorescent cells appeared to be a heterogeneous population, suggesting that multiple PylRS variants enabled the aminoacylation of ncAA **1**.

**Figure 3.**
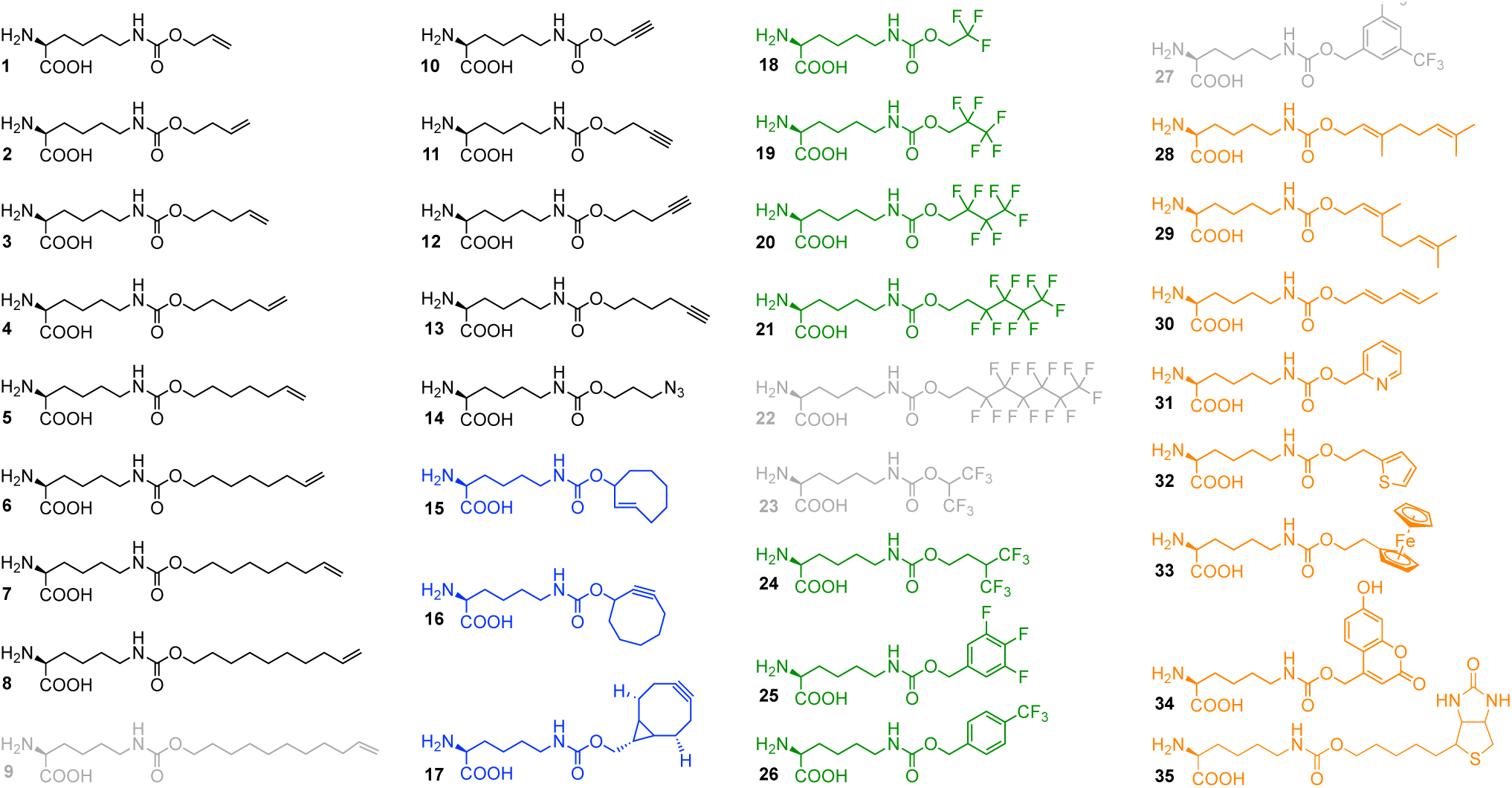
Chemical structures of analyzed ncAAs. Class I: linear alkenes, alkynes, and azide (black). Class II: cyclic alkenes and alkynes (blue). Class III: fluorinated side chains (green). Class IV: diverse functionalities (orange). Grey ncAA structures were rejected.

**Figure 4.**
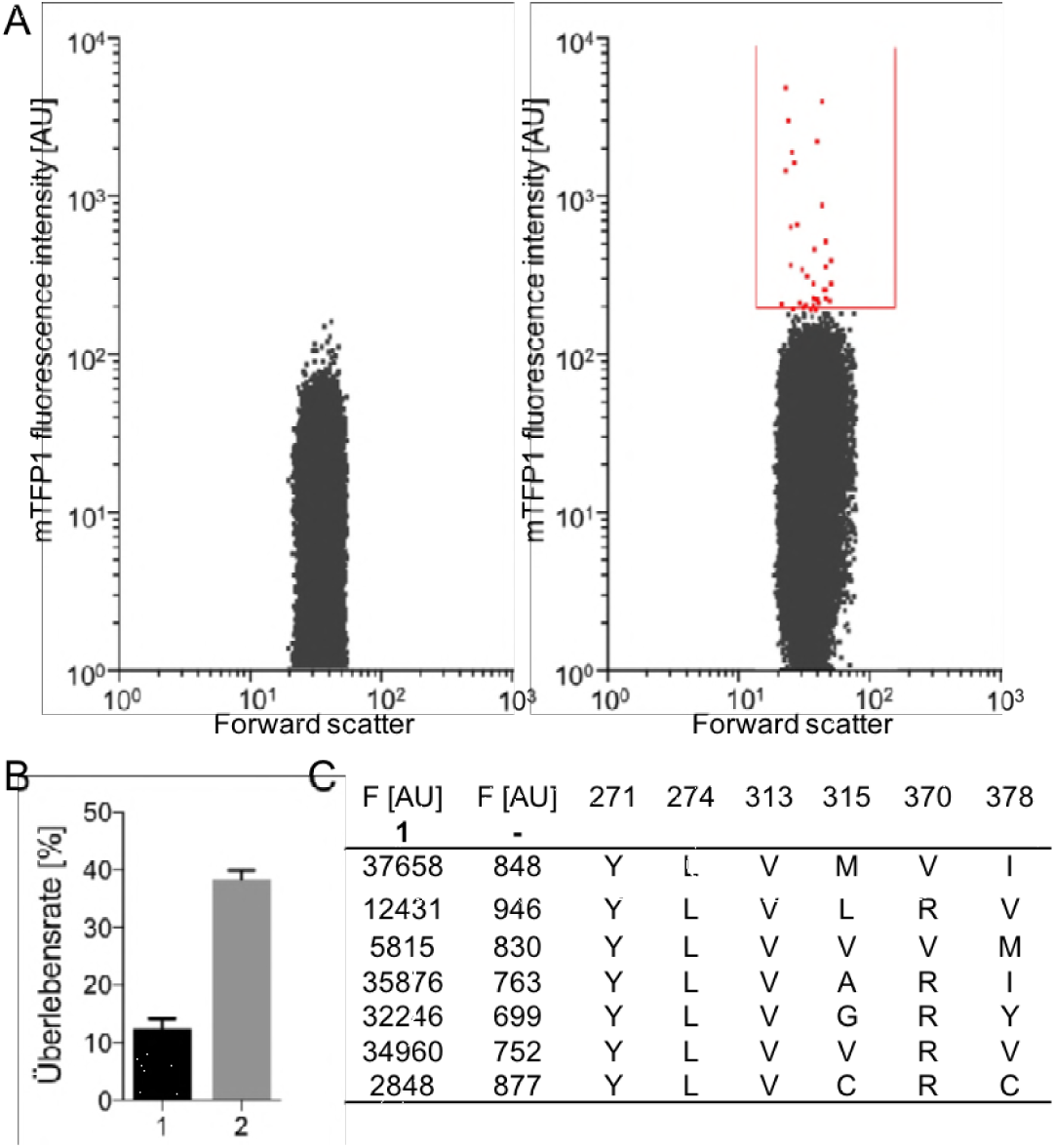
Identification of PylRS variants for the incorporation of ncAA **1**. (A) (Left plot) *E. coli* cells harboring pEVOL303_Lib were grown in the absence of ncAA and analyzed by FACS. Fluorescence intensity in arbitrary units (AU) and forward scatter of each cell were monitored. (Right plot) Cells harboring pEVOL303_Lib were grown in the presence of ncAA **1**. Fluorescent cells (red) within the sort gate (red line) were separated from nonfluorescent cells. (B) Survival rate of cells after FACS in LB medium (1) and conditioned medium (2). Results were obtained from five independent experiments; standard deviation is indicated by error bars. (C) Fluorescence intensities of mTFP1_TAG128_ with **1** and in the absence of ncAA (–) are depicted for seven PylRS variants; the six positions in the mutant library are shown.

All fluorescent cells exceeding the set threshold were sorted on a 96-well plate supplied with conditioned medium and recovered overnight. The medium contained cell-secreted growth factors that improved the survival rate of single cells three-fold compared to the untreated Lysogeny broth (LB) medium (Figure 4B). In the case of ncAA **1**, seven cells were recovered after FACS screening, the plasmid pEVOL303_Lib was purified and the PylRS variants were sequenced (Figure 4C). Next, expression of mTFP1_TAG128_ was repeated in the presence of the PylRS variants, the tRNA^Pyl^ and ncAA **1** in *E. coli* cells. In all cases, the fluorescence measurements were in agreement with the FACS results and did not reveal any background expression of mTFP1 in the absence of ncAA **1** (Figure 4C).

Overall, 151 PylRS variants (110 unique and 41 redundant) facilitating the incorporation of 20 out of the 23 tested ncAAs were developed. Seventeen PylRS variants accepted multiple ncAAs. The ncAAs were aminoacylated by up to 16 different PylRS (Supplementary Table 2). The amino acid abundancies of the six mutated positions (P1–P6) of all sorted PylRS variants are summarized for each ncAA in a heat map (Figure 5).

**Figure 5.**
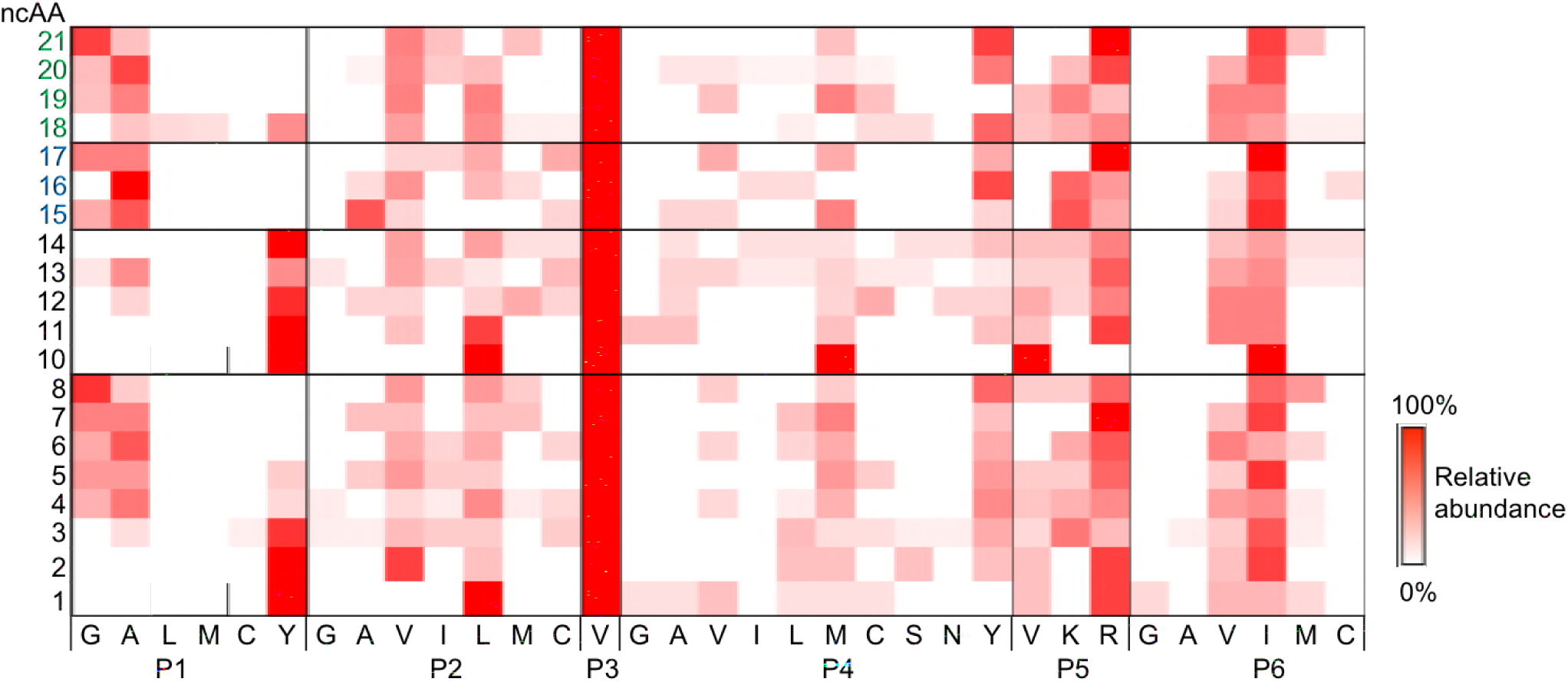
Heat map of the analyzed ncAAs **1-23** versus the six variable positions of the PylRS library (P1=Y271, P2=L274, P3=C313, P4=M315, P5=V370, P6=I378). Amino acids are represented by a one-letter code; the color intensity indicates the relative abundance of a mutation (white: 0%; red: 100%). Modifications not present in any of the sequenced PylRS variants are omitted. The incorporation of ncAAs **9**, **22**, and **23** was not detected.

We divided the tested ncAAs **1–23** into three distinct structural classes (Figure 3). Class I ncAAs **1–14** bear a linear aliphatic tail. Like a molecular ruler, these ncAAs allow to assess the maximum carbon chain length accepted by PylRS based on the fluorescent signal of the mTFP1 reporter. For example, ncAA **9** with 11 carbon atoms following the carbamate function exceeded the permissible chain length. Hence, no incorporation was detected. Interestingly, the cells tended to aggregate upon addition of ncAA **9** (Supplementary Figure 1). Our data suggest that the long hydrophobic tail of ncAA **9** cross-linked the cells and prevented its uptake into the cytoplasm. Class II ncAAs **15–17** are cyclic and can be used in strain-promoted cycloadditions for labeling of proteins (25, 26). The higher steric demand of fluorinated side chains of class III ncAAs **18–23** might single out those PylRS variants that accept particularly bulky ncAAs, since the volume of a CF_3_ substituent (39.8 Å^3^) is about twice as big as a methyl-group (21.6 Å^3^) (27). Interestingly, the FACS screen detected the aminoacylation of the ncAAs **18–21**, but not ncAA **22** or **23** (Figure 5). These findings show that the (CH_2_)_2_(CF_2_)_3_-CF_3_ side chain of ncAA **21** appeared to define the maximum length of fluorinated ncAAs in our PylRS library. Notably, ncAA **23**, which is short and bears a branch directly after the carbamate, was not incorporated either. This indicates that at least one carbon atom located between the carbamate and a fluorinated branch is essential for the aminoacylation.

### Pocket-substrate relationship

The heat map of the PylRS variants was analyzed to determine the key residues in the binding pocket for the aminoacylation of the tested ncAAs **1–23**. The 20 incorporated ncAAs required diverse structural adaptions of the active site, due to the different steric demands and chemistries of their side chains (Figure 5). Y271 (P1) defined the bottom of the binding pocket in PylRS; therefore, this position strongly varied with the ncAA size. While tyrosine was highly conserved at P1 for the short ncAAs **1**, **2**, **10**, **11**, and **14**, alanine was favored by the larger ncAAs **4–7**, **13**, **15–17**, **19**, and **20** in agreement with previous reports (9, 28). Aminoacylation of the longest ncAAs **8** and **21** was preferred by a glycine at P1 of the active site. We observed a similar trend for the position L274 (P2), which constituted the rear end of the binding pocket. While leucine was favored by the small ncAAs **1**, **10**, and **11**, the shorter valine occurred in high abundance for larger ncAAs. Interestingly, the mutation C313V (P3), which was previously shown to significantly improve the aminoacylation of ncAAs (29), was strictly conserved throughout all sequenced variants. M315 (P4) differed widely among the small ncAAs. However, tyrosine was more abundant among the larger ncAAs **8**, **16**, **20**, and **21**. The positions V370 (P5) and I378 (P6) were not influenced by the size or chemistry of the incorporated ncAAs. Arginine and isoleucine appeared with the highest frequency among all sorted variants.

### Design of a promiscuous PylRS variant

Orthogonal aaRS/tRNA pairs are usually engineered for a specific ncAA. In some cases, however, an aaRS/tRNA pair may aminoacylate up to 40 ncAAs with similar structural features (30). To design a promiscuous PylRS variant for the incorporation of large and bulky ncAAs, we determined potential key residues from the heat map (Figure 5). We combined the mutations Y271A, L274V, C313V, M315Y, Y349F, and V370R to yield a highly promiscuous PylRS (HpRS).

To test the substrate scope, the HpRS/tRNA^Pyl^ pair and mTFP1_TAG128_ were expressed with and without the four classes of ncAAs **1–35** (1 mM). The HpRS/tRNA^Pyl^ pair worked orthogonally in *E. coli*; thus, the fluorescence signal of mTFP1_TAG128_ was undetectable in the absence of ncAAs (Figure 6). In contrast, the presence of the ncAAs resulted in a significant fluorescent signal (Figure 6A-C). The incorporation of the ncAAs **1–8**, **10–21**, **24–26**, and **28–35** into mTFP1 was additionally confirmed by SDS-PAGE (Supplementary Figure 2) and ESI-TOF (Supplementary Table 3). Remarkably, HpRS tolerated a broad range of lysine carbamates with diverse features, including aliphatic chains, fluorinated residues, cyclic structures, and the biotinylated ncAA **35**. In agreement with our assumption, HpRS was highly capable of aminoacylating ncAAs with long side chains **3-6** and gave only moderate results for the smallest tested ncAAs **1** and **10**.

**Figure 6.**
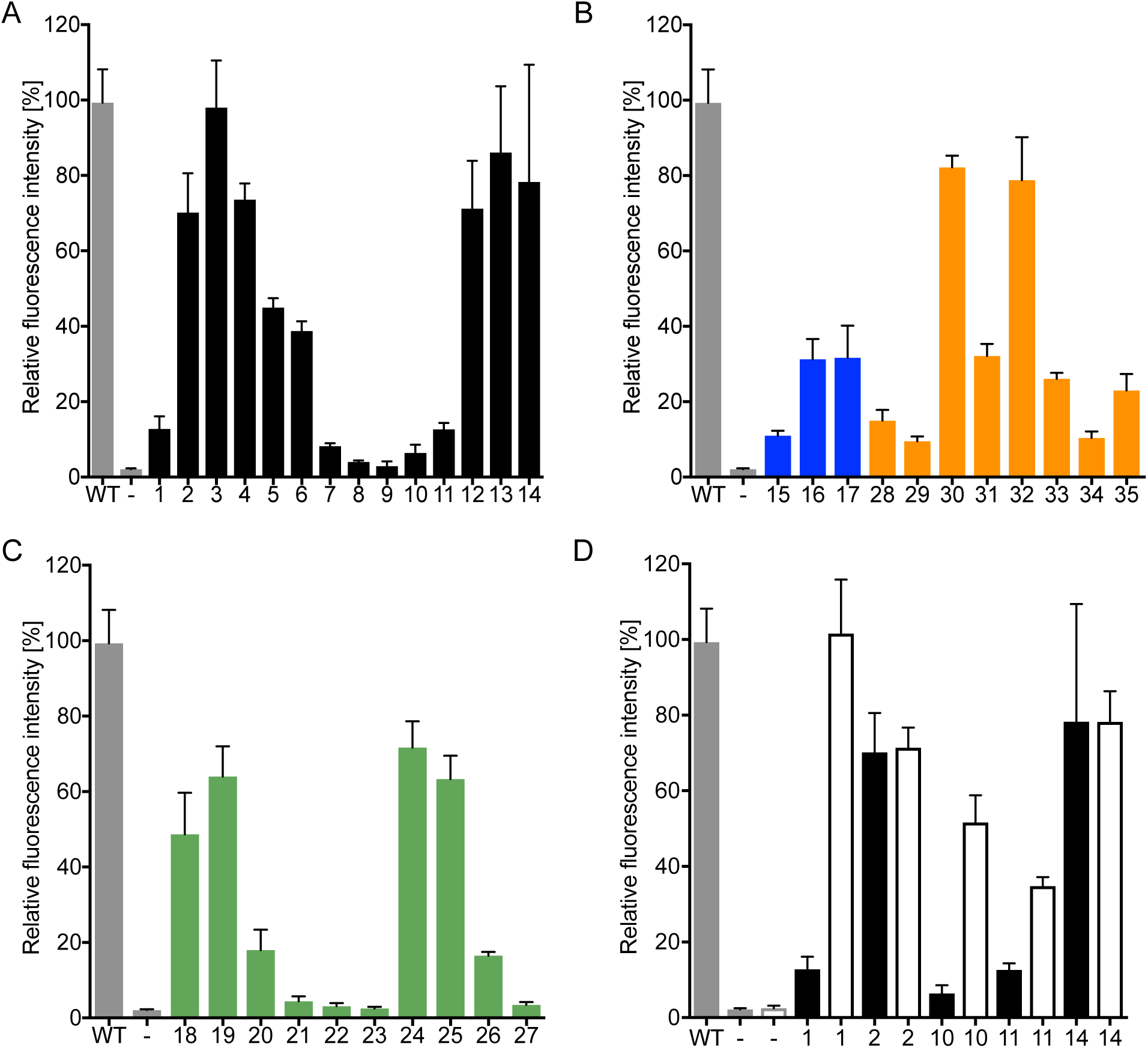
HpRS-dependent incorporation of ncAAs **1–35** in mTFP1_TAG128_. Fluorescence intensity of mTFP1 is normalized against wild-type (WT) mTFP1 (grey bar), while expression of mTFP1_TAG128_ in the absence of ncAAs (**-**, grey bar) served as a negative control. Error bars indicate the standard deviation of three independent experiments. (A) Substrate profiles of HpRS for class I (black): ncAAs **1-14**. (B) Class II (blue): ncAAs **15–17** and class IV (orange): ncAAs **28–35**. (C) Class III (green): ncAAs **18–27**. (D) Comparison of the aminoacylation efficiencies of HpRS (solid) and the PylRS variant (C313V, Y349F) (no fill) for ncAAs **1**, **2**, **10**, **11**, and **14**.

To yield a PylRS variant particular suited for the aminoacylation of small ncAAs, we combined the mutations C313V and Y349F. To validate our heat-map-guided design, we expressed the PylRS variant (C313V, Y349F) together with tRNA^Pyl^ and mTFP_TAG128_ in the presence of the ncAAs **1**, **2**, **10**, **11**, and **14**. Indeed, this PylRS variant (C313V, Y349F) showed a significantly higher aminoacylation efficiency of the small ncAAs **1**, **10**, and **11** than HpRS (Figure 6D). The homology models provide further insights into the structural basis for the observed aminoacylation efficiency (Figure 7). Among the six mutations in HpRS, Y271A had the most pronounced effect on the shape of the binding pocket, opening up a large cavity at the bottom. L274V generated additional spaces. Overall, the binding pocket of HpRS appeared to be substantially different in terms of volume from the PylRS variant (C313V, Y349F).

**Figure 7.**
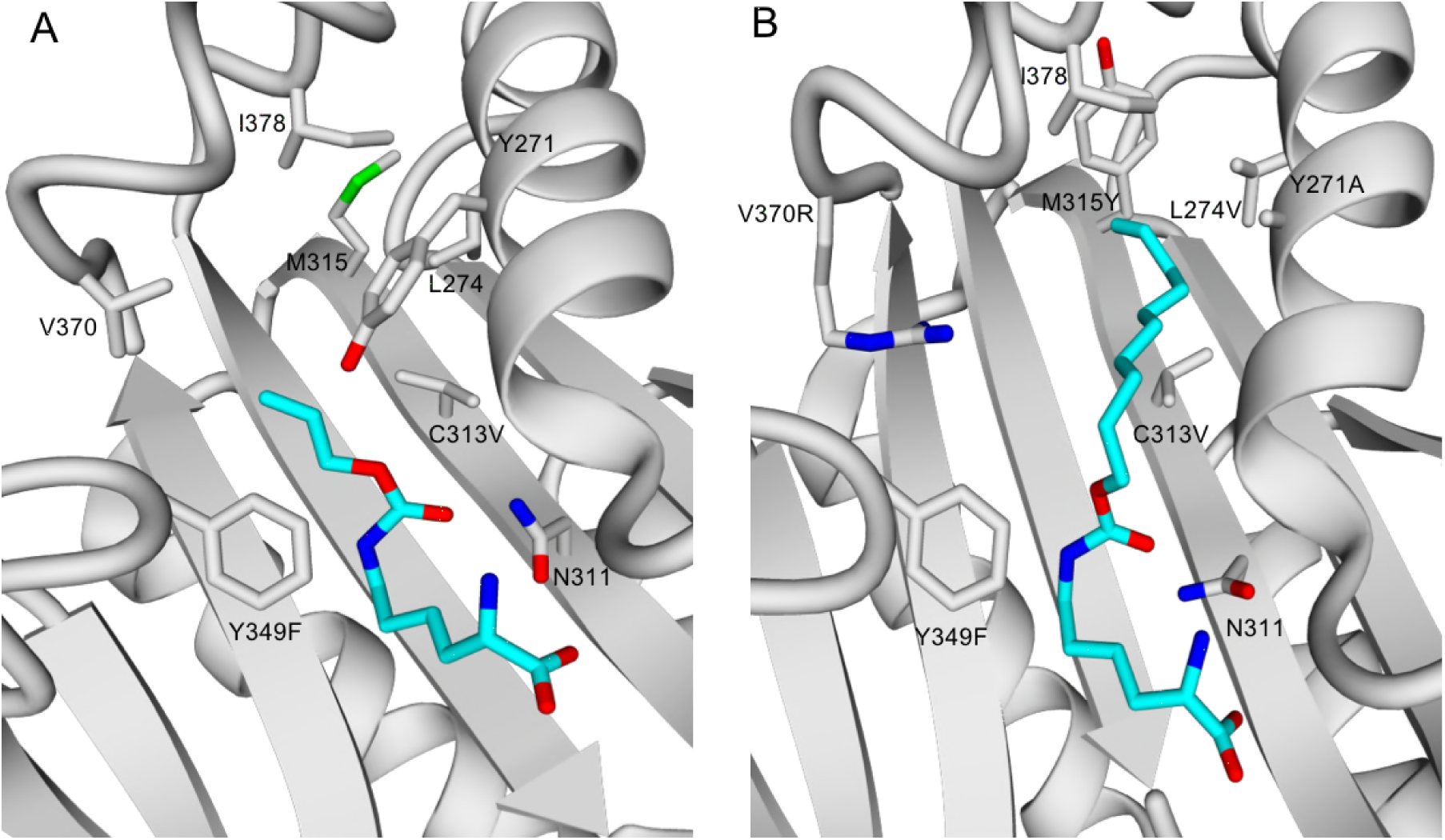
Comparison between the active sites in the PylRS variant (C313V, Y349F) and HpRS. (A) Homology model of the *M. barkeri* PylRS variant (C313V, Y349F) in complex with the ncAA **1** (cyan). (B) Homology model of the *M. barkeri* HpRS in complex with the ncAA **21** (cyan).

## Discussion

The engineering process of ncAA-specific aaRS/tRNA pairs is typically sophisticated and time consuming. In general, directed approaches begin with the composition of a large aaRS mutant library containing 10^8^–10^9^ variants and a subsequent screening to identify the beneficial variants (31). Therefore, dead-and-alive assays on a selective solid or liquid medium are the preferred method for screening the large quantities of variants in a short time as demonstrated on existing aaRS/tRNA pairs (2, 13). Nevertheless, this screening method suffers from a harsh selection pressure that is difficult to adjust, causing slow cell growth and low survival rates (17). Consequently, even beneficial variants might be unintentionally excluded from the library. Additionally, large libraries that likely exceed the transformation efficiency of expression strains, lead to less diversity.

In parallel, more focused libraries have shown to sustain a higher frequency of beneficial variants and to provide promising results in directed evolution (32). Unlike dead-and-alive assays, FACS provides a tunable sorting stringency over a broad dynamic range for the ncAA incorporation (16, 17, 33).

Our FACS-based screen takes advantage of the unique properties of the PylRS, improving speed, sensitivity and selectivity. The position N311 was used as the selection filter instead of iterative rounds of negative and positive selections. The efficiency of our method resulted in a higher diversity of positive mutants that might have been rejected by classical assays. The possibility of a random mutation within the amber codon, which could cause a false positive signal, was reduced due to the smaller number of generations. We maintained the complexity of our mutant library during the FACS screen by using a conditioned medium, which improved the survival rate of the cells three-fold. With our setup, we could demonstrate that a relatively small PylRS library, comprised of six positions (9.5x10^4^ variants), and a single positive selection round of FACS were sufficient for identifying 110 orthogonal PylRS variants that together incorporate 20 structurally diverse ncAAs (Supplementary Table 2).

Our newly established procedure enabeled the prediction of the promiscuous PylRS variant, HpRS, with a particular preference for larger ncAAs. HpRS accepts a broad range of ncAAs (31 out of 35) making this PylRS a valuable tool for biotechnology with multiple applications and an ideal starting point for future mutant libraries. Furthermore, the site-directed incorporation of the biotinylated ncAA **35** into a protein of interest might be a popular starting point to simplify techniques such as Western blotting, ELISA, flow cytometry or antibody labeling kits.

In summary, we demonstrated the ability to rapidly identify beneficial PylRS variants from a focused mutant library for the incorporation of a broad range of ncAAs. The high efficiency of our FACS-based screening method reduced the frequency of false positive signals and facilitated the analysis of the sequence space in the sorted PylRS variants. Our approach enabled the identification of the promiscuous variant, HpRS. Hence, the prediction of PylRS/tRNA^Pyl^ pairs for ncAAs based on the pocket-substrate information can now be used as an example for others to avoid the cumbersome engineering process.

## Material and methods

### Strains

All cloning steps were performed in TOP10 *E. coli* (Thermo Fisher Scientific). The FACS and protein expression experiments were carried out with EXPRESS BL21(DE3) (Lucigen, Middleton, WI).

### Medium

A conditioned medium was used to improve the survival of sorted cells after FACS. To prepare the conditioned medium, BL21 (DE3) cultures were grown in LB (0D_600_~1.0), centrifuged at 8000 x g for 20 min at 4 °C and the supernatant was sterilized by passing through a 0.22 μm cellulose filter (Millipore, Bedford, MA).

### Non-canonical amino acids (ncAA)

ncAA **1** (Sigma-Aldrich), ncAA **15–17** (Sirius Fine Chemicals, Bremen, Germany), and ncAAs **24–27**, **30–33**, as well as **35** (SUNGYOUNG Chemical Limited, Shanghai, China) were purchased from commercial sources. ncAA **34** was synthesized according to the procedure published by Luo *et al.* (34). ncAA **2–14**, **18–23**, **28**, and **29** were synthesized according to a modified procedure from Li *et al.* (35) (Supplementary Information (S2)).

### Construction of pEVOL303

The plasmid pEVOL303 was constructed by ligating the plasmid pEVOL (36) with the commercially available pET303/CT-His (Thermo Fisher Scientific) using Gibson cloning (37). Prior to ligation, pEVOL was digested with the restriction enzymes SacI and *PciI*. The required region of pET303 was amplified using the primers pET303_f and pET303_r (Supplementary Table 1). The resulting plasmid, pEVOL303, carried a p15A origin, a chloramphenicol resistance gene, a *Methanosarcina barkeri* PylRS gene downstream of a pBAD promoter, and the corresponding *tRNA* gene controlled by a proK promoter. The fluorescent protein gene with an N-terminal 6xHis-SUMO tag, either wild-type mTFP1 or mTFP1_TAG128_ (bearing an amber codon at position 128), was cloned downstream of a T7 promoter. The amber codon was inserted using the primers mTFP_128_f and mTFP_128_r (Supplementary Table 1).

### PylRS library construction

*Methanosarcina barkeri* PylRS containing the single mutation Y349F served as the starting point for our mutant library. Six positions in PylRS, Y271, L274, C313, M315, V370, and I378, were selected for our library based on a homology model derived from the crystal structure of PylRS (PDB 4Q6G and 4CS3) (Figure 1A and B) (38, 39). A pre-defined set of amino acids was inserted into the six positions, resulting in 95,040 mutants. To ensure an equal distribution of all the mutants, the library was assembled commercially by the company, Life Technologies (Thermo Fisher Scientific) and introduced to pEVOL303 *via* the restriction sites *SalI* and *BglII*. The resulting plasmid, pEVOL303_Lib, was transformed into BL21 (DE3) cells applying the standard electroporation protocol. The transformation was determined with 1×10^7^ CFU per preparation.

### Sample preparation for FACS

BL21 (DE3) cells transformed with pEVOL303_Lib were inoculated in 50 mL LB medium supplemented with 25 μg/ml chloramphenicol (Cm), and grown until an OD_600_ of 0.7. Then, 450 μL of the cells was transferred into a 2 mL reaction vessel and induced with 50 μL induction medium (10 mM ncAA, 10 mM IPTG, 1% w/v arabinose) and incubated for 16 h at 37 °C and 700 rpm. The cells were diluted to a final concentration of 1×10^7^ cell/mL and washed twice with an M9 minimal medium supplemented with 25 μg/ml chloramphenicol. In the beginning of each FACS experiment, the negative control (cell growth in the absence of ncAA) was screened first to ensure the discrimination of canonical amino acids by the PylRS mutant library. Cell sorting was performed by detecting the fluorescence of mTFP1 with a BD Influx (BD Biosciences) operated with filter-sterilized BD FACS Flow Sheath Fluid, a 457 nm laser for excitation, and a 480/40 bandpass filter. The selected operation mode was 1.0 Drop Single. The selection threshold (gate) was adjusted based on the fluorescent signal from the first 1×10^5^ cells of each sample. The fluorescence and side scatter of 1×10^7^ cells were monitored. Cells within the gate were sorted separately on a 96-well plate (Sigma-Aldrich) supplied with the conditioned medium containing 25 μg/ml chloramphenicol. The growth of the sorted cells was continued overnight at 37 °C and 300 rpm. Each culture was sequenced using the primers Lib_seq_f and Lib_seq_r (Supplementary Table 1). pEVOL303_Lib containing the sequenced PylRS variants were retransformed into new BL21 (DE3) cells, grown and induced with the induction medium at OD_600_ of 0.7. Correspondingly, PylRS, tRNA^Pyl^, and mTFP1_TAG128_ were expressed. Both the incorporation of the ncAAs and the orthogonality of the PylRS variant were confirmed by fluorescence measurement of mTFP1 (excitation 462 nm, emission 492 nm) in a black, 96-well plate (Thermo Fisher Scientific) with an Infinite M1000 plate reader (Tecan, Zurich, Switzerland).

### ncAA incorporation by HpRS and Variant (C313V and Y349F)

A single BL21 (DE3) colony containing pEVOL303 encoding for HpRS or PylRS (C313V and Y349F) was selected from the LB/Cm agar plate and inoculated into 50 mL LB/Cm media, then incubated and shaken overnight at 37 °C. The resulting culture was diluted (1:100) with fresh LB/Cam media and grown to an OD_600_ of 1.5. The mTFP1 expression was triggered by adding 1 mM ncAA, 1 mM IPTG, and 0.1% w/v arabinose and cell growth continued at 37 °C and 700 rpm for 16 h. Fluorescence was measured correspondingly. In the end, the sample was heated at 75 °C for 20 min and centrifuged at 20,000 × g for 15 min. The supernatant was analyzed by SDS-PAGE.

### Mass spectrometry

Harvested *E. coli* Bl21 (DE3) cells were washed with phosphate-buffered saline (PBS). The cell pellet was suspended in lysis buffer (100 mM Tris pH 7.5, 500 mM NaCl, 20 mM imidazole, 10%, v/v glycerol) and incubated at 75 °C for 20 min. Cell debris was removed by centrifugation (60,000 x g, 30 min at 4 °C). The supernatant was loaded onto a HisTrap HP Ni–NTA column (GE Healthcare), pre-equilibrated with lysis buffer. The mTFP1 was eluted by increasing the concentration of imidazole from 20 mM to 500 mM in the lysis buffer over 10 column volumes. SUMO protease was added to the eluted protein, dialyzed overnight at 4 °C against ddH_2_O and passed through a HisTrap HP Ni–NTA to remove the remaining N-terminal 6xHis as well as the SUMO tag. Then mTFP1 was incubated at 75 °C for 10 min, centrifuged at 60,000 × g for 30 min at 4 °C and analyzed by mass spectrometry (maXis HD^TM^ ESI-TOF, Bruker) at a concentration of 0,04 mM. The sample was injected into a high-performance liquid chromatography (Agilent Technologies, C4 column, column volume of 5 mL) and separation was performed at a constant flow rate of 0.5 μL and a gradient of 80% acetonitrile and 0.1% formic acid for 8 min. Fractions were recorded according to the standard procedure.

### Homology model of the *Methanosarcina barkeri* PylRS in complex with pyrrolysine

Homology models were generated using the YASARA Structure, Version 14.7.17 (40). The catalytic domain of *Methanosarcina barkeri* PylRS served as the template for YASARA’s homology modeling macro, using the conservative “slow” protocol with the following parameter settings: number of PSI-BLAST iterations – 10; maximum allowed BLAST E-value to consider template – 0.5; maximum oligomerization state – 2; maximum number of alignment variations per template – 4; maximum number of conformations tried per loop – 200; and maximum number of residues added to the termini – 20. The resulting homology model was based on two structures of PylRS variants, PDB 4Q6G and 4CS3; both sequences had a 98% homology with the template (38, 39). The YASARA algorithm performed the secondary structure prediction, loop construction, and amino-acid rotamer selection, followed by a steepest-descent energy minimization.

### Model of *Methanosarcina barkeri* PylRS mutants in complex with ncAAs

To create HpRS and PylRS (C313V, Y349F), mutations were introduced to the PylRS homology model. Next, the bound pyrrolysine ligand was converted into the designated ncAA *in silico*. Subsequently, all of the residues in the model, except for the mutation sites and the ncAA side chain, were frozen for a molecular-dynamics simulation (50 ps, T = 300 K) with YASARA’s recommended default force field, AMBER 2003 (41). Eventually, the energy of the free residues was repeatedly minimized with the AMBER 2003 force field until overlays with the previous model did not reveal any significant changes in the orientation of the respective side chains.

## Acknowledgements

The research reported in this publication was supported by funding from King Abdullah University of Science and Technology (KAUST). We thank the SFB749/A10 (M.G.) for financial support. We are grateful to Prof. Peter G. Schultz (The Scripps Research Institute, La Jolla, CA) for kindly providing the original pEVOL-PylRS plasmid.

## Author contributions

J. E., M. G., M. R. designed and supervised the research project; A. H., D. R. and A. A. performed the molecular biology; A. H. and A. D. conducted the FACS experiments; X. L. and S. G. synthesized the ncAAs; J. E., A. A., R. K., M. R., and A. H. analyzed the data and wrote the manuscript.

## S1. Additional Tables and Figures

**Supplementary Table 1.**
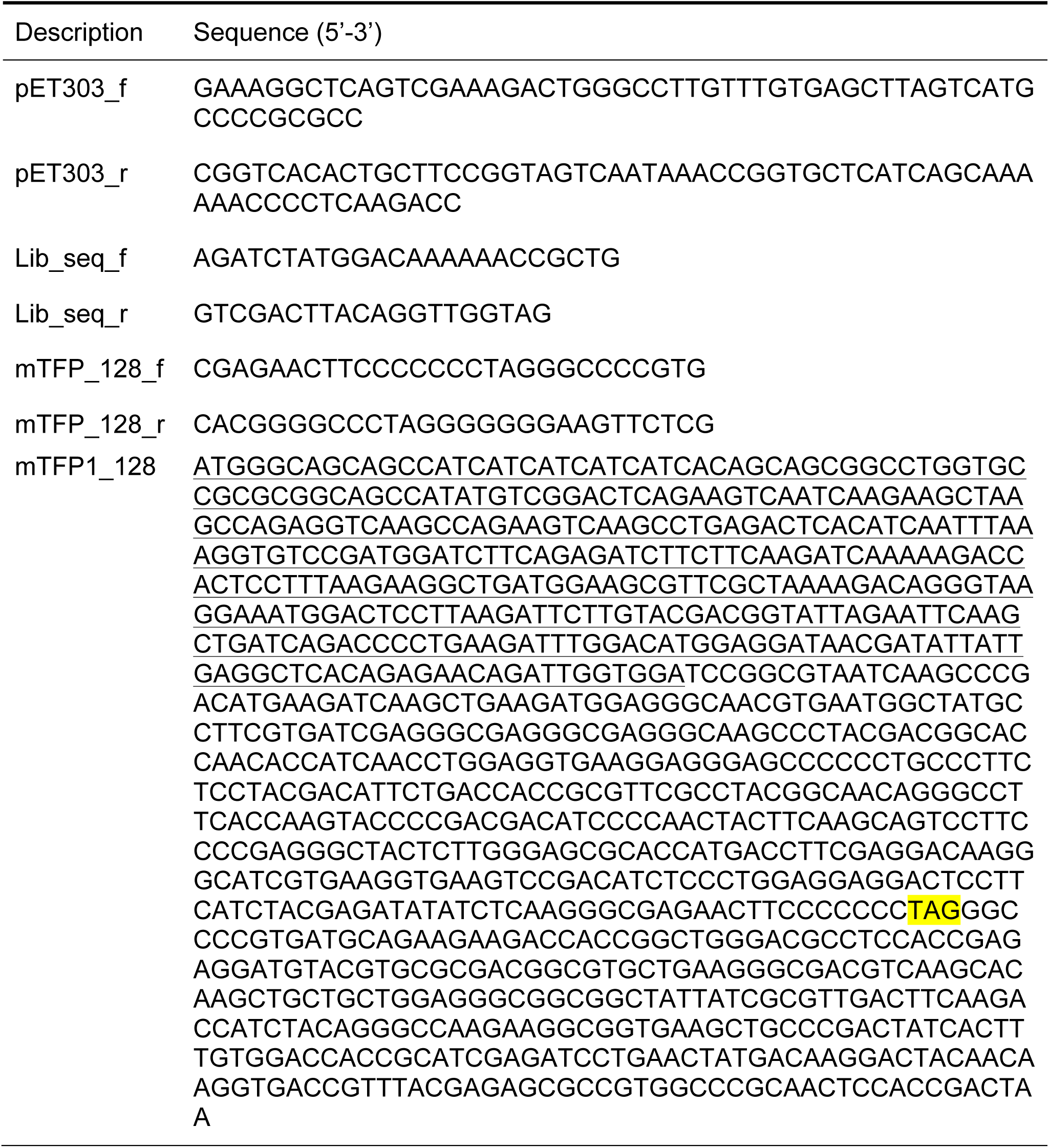
List of oligomers used in this study. Amber codon is highlighted yellow. Underscored sequence encodes the 6xHis-Sumo tag.

**Supplementary Table 2.** List of 110 unique PylRS variants and their corresponding substrate scopes with respect to the ncAAs. A total of 151 PylRS variants were sequenced. Only the six positions in PylRS that were included in the mutation library are depicted. The normalized fluorescence intensity of mTFP1_TAG128_ (F [%]) co-expressed with a PylRS/tRNA^Pyl^ pair, is listed both with (+) and without (-) ncAAs (1 mM). Fluorescence intensity was normalized against wild-type mTFP1. PylRS variants were not detected for ncAAs **9, 22, or 23**.

| Variant | Position |  |  |  |  |  | ncAA | F [%] + | F [%] - |
| --- | --- | --- | --- | --- | --- | --- | --- | --- | --- |
|  | P1<br>271 | P2<br>274 | P3<br>313 | P4<br>315 | P5<br>370 | P6<br>378 |  |  |  |
| 1 | Y | L | V | M | V | I | 1 | 94 | 2 |
|  |  |  |  |  |  |  | 4 | 33 | 2 |
|  |  |  |  |  |  |  | 10 | 48 | 1 |
|  |  |  |  |  |  |  | 12 | 35 | 2 |
|  |  |  |  |  |  |  | 13 | 23 | 2 |
|  |  |  |  |  |  |  | 14 | 87 | 1 |
| 2 | Y | L | V | L | R | V | 1 | 31 | 2 |
| 3 | Y | L | V | V | V | M | 1 | 15 | 2 |
| 4 | Y | L | V | A | R | I | 1 | 90 | 2 |
| 5 | Y | L | V | G | R | Y | 1 | 81 | 2 |
| 6 | Y | L | V | V | R | V | 1 | 87 | 2 |
| 7 | Y | L | V | C | R | C | 1 | 7 | 2 |
| 8 | Y | V | V | S | R | I | 2 | 32 | 2 |
| 9 | Y | V | V | L | R | V | 2 | 27 | 1 |
| 10 | Y | V | V | M | V | I | 2 | 68 | 1 |
| 11 | Y | V | V | Y | R | I | 2 | 32 | 1 |
|  |  |  |  |  |  |  | 11 | 90 | 2 |
| 12 | Y | A | V | N | K | A | 3 | 3 | 2 |
| 13 | C | L | V | L | R | I | 3 | 11 | 2 |
| 14 | Y | V | V | S | K | I | 3 | 45 | 2 |
| 15 | Y | I | V | L | K | I | 3 | 42 | 2 |
| 16 | Y | C | V | C | K | V | 3 | 75 | 2 |
| 17 | A | L | V | Y | K | M | 3 | 22 | 2 |
|  |  |  |  |  |  |  | 18 | 45 | 2 |
| 18 | Y | C | V | L | K | I | 3 | 75 | 2 |
| 19 | Y | V | V | Y | K | I | 3 | 68 | 2 |
| 20 | Y | G | V | Y | R | V | 3 | 27 | 2 |
| 21 | Y | V | V | M | R | I | 3 | 71 | 2 |
| 22 | A | C | V | L | V | I | 3 | 26 | 2 |
| 23 | Y | I | V | Y | K | I | 3 | 70 | 2 |
| 24 | Y | I | V | Y | V | I | 3 | 78 | 2 |
| 25 | Y | V | V | C | R | V | 3<br>18 | 74<br>38 | 2<br>2 |
| 26 | A | V | V | Y | R | I | 3<br>5<br>7<br>16<br>18<br>20<br>20<br>21 | 108<br>44<br>13<br>33<br>57<br>21<br>17<br>5 | 2<br>2<br>2<br>2<br>2<br>2<br>2<br>2 |
| 27 | A | I | V | V | R | V | 4<br>13<br>20 | 56<br>5<br>7 | 2<br>1<br>2 |
| 28 | Y | C | V | M | V | V | 4 | 90 | 2 |
| 29 | G | L | V | M | K | V | 4 | 61 | 2 |
| 30 | A | L | V | Y | K | V | 4<br>16<br>20 | 71<br>22<br>11 | 2<br>2<br>2 |
| 31 | A | L | V | Y | R | I | 4 | 77 | 3 |
| 32 | G | L | V | Y | R | I | 4<br>17 | 54<br>22 | 2<br>2 |
| 33 | Y | G | V | V | R | I | 4 | 79 | 2 |
| 34 | A | V | V | L | R | I | 4<br>6<br>7<br>8<br>20 | 70<br>6<br>6<br>5<br>6 | 2<br>2<br>2<br>2<br>2 |
| 35 | G | M | V | M | V | I | 4 | 45 | 2 |
| 36 | A | V | V | M | R | I | 4 | 78 | 2 |
| 37 | A | C | V | Y | K | M | 4<br>13 | 68<br>64 | 2<br>2 |
| 38 | A | L | V | Y | V | V | 4 | 67 | 2 |
| 39 | G | L | V | M | R | I | 5<br>7<br>8<br>17<br>17 | 49<br>8<br>4<br>32<br>33 | 2<br>2<br>2<br>2<br>2 |
| 40 | Y | A | V | C | V | V | 5 | 28 | 2 |
| 41 | G | V | V | M | R | I | 5 | 36 | 2 |
| 42 | A | I | V | Y | K | I | 5<br>20 | 25<br>10 | 2<br>2 |
| 43 | G | L | V | V | R | V | 6 | 29 | 2 |
| 44 | A | L | V | M | K | M | 6 | 17 | 2 |
| 45 | A | C | V | Y | R | V | 6 | 33 | 2 |
| 46 | A | I | V | M | K | V | 6 | 21 | 2 |
| 47 | G | V | V | Y | R | I | 6<br>8<br>20<br>20<br>20<br>21<br>21 | 37<br>4<br>19<br>24<br>14<br>3<br>3 | 2<br>2<br>2<br>2<br>2<br>2<br>2 |
| 48 | G | M | V | M | R | I | 7<br>8<br>21 | 5<br>4<br>5 | 2<br>2<br>2 |
| 49 | A | A | V | Y | R | V | 7<br>8 | 5<br>9 | 2<br>2 |
| 50 | G | M | V | M | R | V | 7 | 5 | 2 |
| 51 | Y | L | V | M | R | I | 11 | 38 | 2 |
| 52 | Y | L | V | A | R | V | 11 | 39 | 2 |
| 53 | Y | L | V | G | R | V | 11 | 13 | 2 |
| 54 | Y | C | V | C | V | V | 12 | 40 | 2 |
| 55 | Y | M | V | C | K | V | 12 | 11 | 2 |
| 56 | Y | V | V | N | R | I | 12 | 10 | 2 |
| 57 | A | M | V | Y | R | V | 12 | 23 | 2 |
| 58 | Y | A | V | A | R | I | 12 | 24 | 2 |
| 59 | A | I | V | M | R | V | 13 | 32 | 2 |
| 60 | Y | Y | V | V | R | V | 13 | 23 | 2 |
| 61 | Y | V | V | A | R | C | 13 | 30 | 2 |
| 62 | Y | C | V | S | R | I | 13 | 40 | 2 |
| 63 | G | V | V | L | V | I | 13 | 16 | 2 |
| 64 | Y | G | V | A | R | V | 13 | 69 | 2 |
| 65 | A | C | V | I | R | I | 13 | 53 | 2 |
| 66 | A | V | V | C | K | I | 13<br>20 | 45<br>6 | 2<br>2 |
| 67 | Y | V | V | L | R | I | 14 | 87 | 1 |
| 68 | Y | L | V | I | R | I | 14 | 81 | 2 |
| 69 | Y | M | V | N | R | V | 14 | 57 | 2 |
| 70 | Y | V | V | A | V | M | 14 | 74 | 2 |
| 71 | Y | V | V | Y | K | M | 14 | 73 | 2 |
| 72 | Y | C | V | S | R | C | 14 | 31 | 2 |
| 73 | Y | L | V | Y | K | V | 14 | 26 | 1 |
| 74 | G | C | V | Y | K | I | 15 | 7 | 2 |
| 75 | A | A | V | V | K | I | 15 | 7 | 2 |
| 76 | A | A | V | L | K | I | 15 | 6 | 2 |
| 77 | G | A | V | L | R | I | 15 | 8 | 2 |
| 78 | A | A | V | L | R | V | 15 | 8 | 2 |
| 79 | A | V | V | A | K | I | 15 | 8 | 2 |
| 80 | A | A | V | I | K | I | 16 | 10 | 2 |
| 81 | A | L | V | Y | K | I | 16 | 16 | 2 |
|  |  |  |  |  |  |  | 18 | 29 | 2 |
| 82 | A | V | V | L | K | I | 16 | 13 | 2 |
| 83 | A | V | V | Y | R | C | 16 | 25 | 2 |
| 84 | A | M | V | Y | R | I | 16 | 11 | 2 |
| 85 | A | C | V | Y | R | I | 17 | 34 | 2 |
| 86 | A | C | V | V | R | I | 17 | 28 | 2 |
| 87 | G | I | V | C | R | I | 17 | 27 | 2 |
| 88 | M | L | V | Y | R | I | 18 | 36 | 2 |
| 89 | Y | C | V | Y | R | C | 18 | 32 | 2 |
| 90 | L | M | V | Y | R | I | 18 | 29 | 2 |
| 91 | Y | V | V | V | R | V | 18 | 37 | 2 |
| 92 | F | V | V | S | V | V | 18 | 4 | 2 |
| 93 | Y | L | V | L | K | V | 18 | 65 | 2 |
| 94 | Y | V | V | Y | K | V | 18 | 25 | 2 |
| 95 | M | L | V | Y | V | I | 18 | 30 | 2 |
| 96 | Y | L | V | S | R | V | 18 | 47 | 2 |
| 97 | A | L | V | V | K | V | 19 | 45 | 2 |
| 98 | Y | V | V | C | R | I | 19 | 25 | 2 |
| 99 | G | L | V | M | V | I | 19 | 53 | 2 |
| 100 | A | L | V | M | K | V | 19 | 63 | 2 |
| 101 | A | L | V | A | R | I | 20 | 6 | 2 |
| 102 | A | I | V | A | R | I | 20 | 7 | 2 |
| 103 | A | L | V | Y | R | V | 20 | 24 | 2 |
| 104 | G | L | V | M | K | I | 20 | 12 | 2 |
| 105 | G | I | V | Y | R | V | 20 | 5 | 2 |
| 106 | A | L | V | M | R | V | 20 | 11 | 2 |
| 107 | A | V | V | V | K | I | 20 | 8 | 2 |
| 108 | A | A | V | I | R | I | 20 | 5 | 2 |
| 109 | A | V | V | Y | R | V | 20 | 12 | 2 |
| 110 | A | I | V | Y | R | M | 21 | 4 | 2 |

**Supplementary Table 3.** ESI-TOF data from purified wild-type mTFP1 (WT) and mTFP1 containing ncAAs **1–35**.

| ncAA | Theoretical Mass (Da) | Experimental Mass (Da) | ± Mass (Da) |
| --- | --- | --- | --- |
| WT<sup>a</sup> | 25154.5 | 25154.7 | 0.2 |
| 1 | 25252.6 | 25251.8 | 0.8 |
| 2 | 25266.6 | 25265.1 | 1.5 |
| 3 | 25280.7 | 25280.2 | 0.5 |
| 4 | 25294.7 | 25294.4 | 0.3 |
| 5 | 25308.7 | 25306.2 | 2.5 |
| 6 | 25322.7 | 25320.9 | 1.8 |
| 7 | 25336.8 | 25334.9 | 1.9 |
| 8 | 25350.8 | 25349.3 | 1.5 |
| 9 | 25364.8 | NA | NA |
| 10 | 25250.6 | 25250.2 | 0.4 |
| 11 | 25264.6 | 25264.4 | 0.2 |
| 12 | 25278.6 | 25277.3 | 1.3 |
| 13 | 25292.7 | 25291.4 | 1.3 |
| 14 | 25295.6 | 25293.4 | 2.2 |
| 15 | 25320.7 | 25318.9 | 1.8 |
| 16 | 25318.7 | 25318.5 | 0.2 |
| 17 | 25344.7 | 25342.3 | 2.4 |
| 18 | 25294.6 | 25293.1 | 1.5 |
| 19 | 25344.6 | 25342.2 | 2.4 |
| 20 | 25394.6 | 25394.6 | 0.0 |
| 21 | 25458.6 | 25457.1 | 1.5 |
| 22 | 25558.6 | NA | NA |
| 23 | 25362.6 | NA | NA |
| 24 | 25390.6 | 25389.1 | 1.5 |
| 25 | 25356.6 | 25355.5 | 1.1 |
| 26 | 25370.7 | 25368.5 | 2.2 |
| 27 | 25438.7 | NA | NA |
| 28 | 25348.8 | 25347.8 | 1.0 |
| 29 | 25348.8 | 25347.5 | 1.3 |
| 30 | 25292.7 | 25290.5 | 2.2 |
| 31 | 25303.6 | 25302.1 | 1.5 |
| 32 | 25322.7 | 25321.4 | 1.3 |
| 33 | 25438.6 | 25437.1 | 1.5 |
| 34 | 25386.7 | 25384.6 | 2.1 |
| 35 | 25424.8 | 25423.9 | 0.9 |

a) WT has the amino acid asparagine (Asn) in position 128.

**Supplementary Figure 1.**
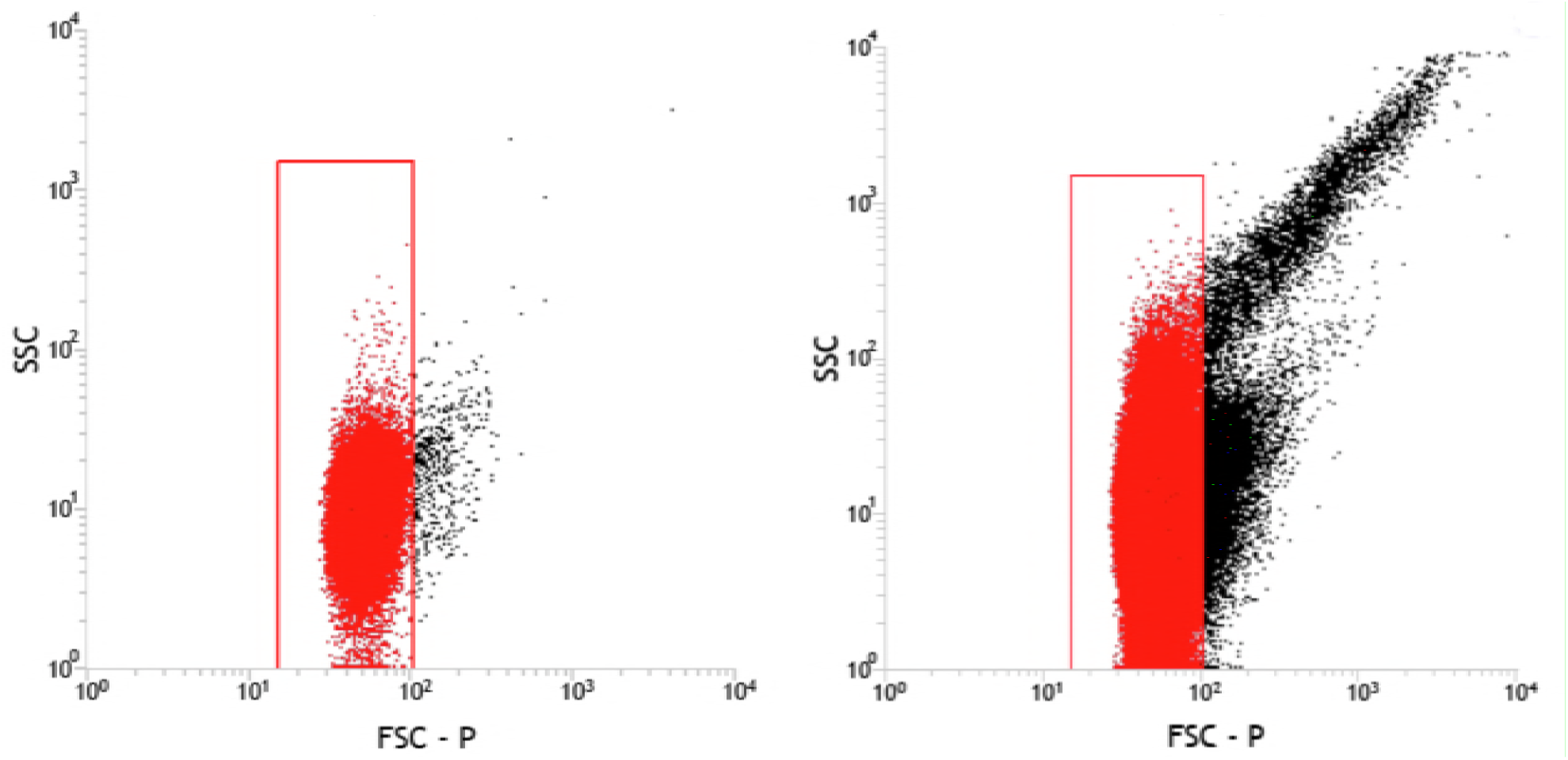
*E. coli* cells harboring pEVOL303_Lib were grown with either ncAA **1** (left) or ncAA **9** (right). The forward (FSC-P) and side (SSC) scatters of each cell were analyzed by FACS. Cells located inside the sorting gate are indicated by the red box. The increased forward and side scatter signals upon ncAA **9** addition indicated cell aggregation.

**Supplementary Figure 2.**
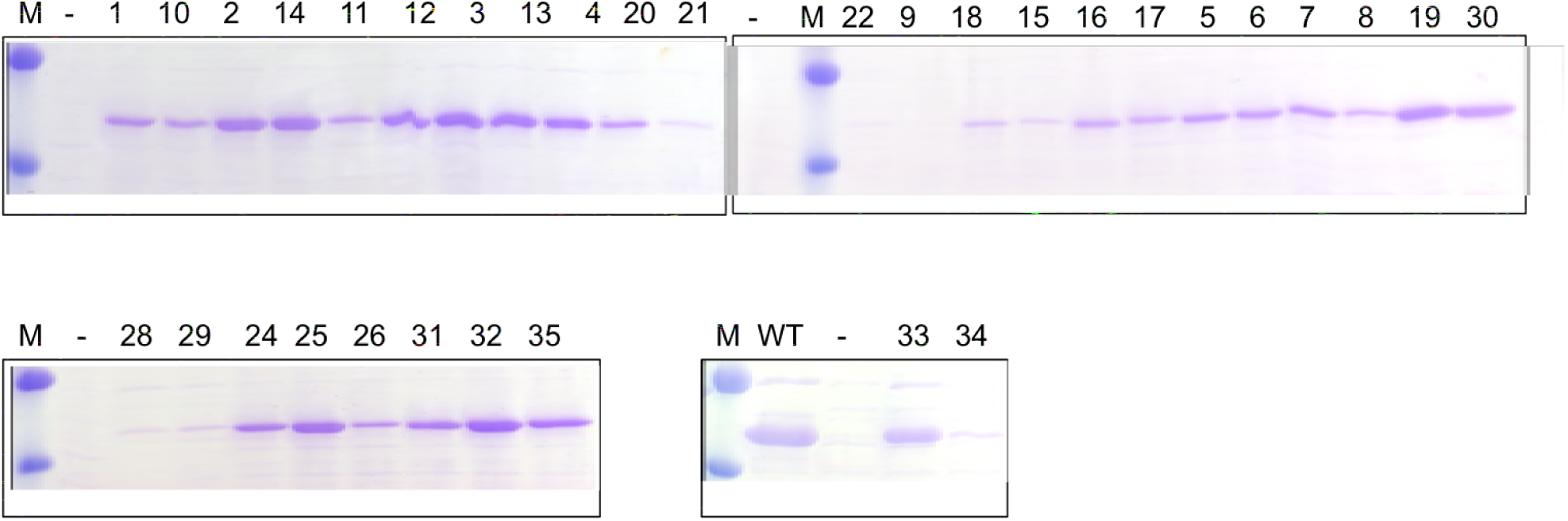
SDS-PAGE analysis of the heat-treated *E. coli* lysate containing mTFP1 with an N-terminal His_6_-SUMO tag. mTFP1_TAG_128 was co-expressed with the HpRS/tRNA^Pyl^ pair, both with and without ncAAs **1–35** (1 mM). ncAA **23** and **27** were not incorporated into the mTFP1 (data not shown). Marker bands indicate 36 and 55 kDa.

## S2. Synthesis

### S2.1 General Procedure

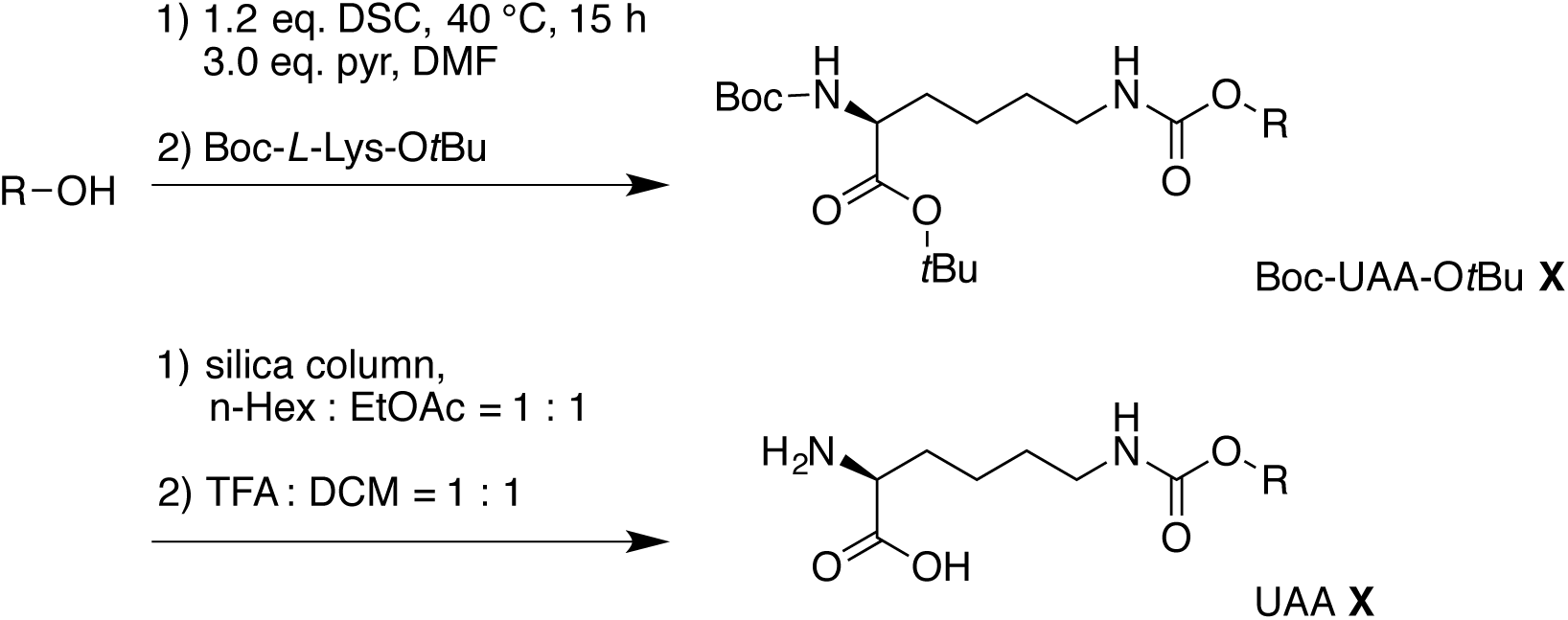

N,N-Disuccinimidyl carbonate (DSC) (0.613 g, 1.2 eq.) and pyridine (0.28 ml, 3.0 eq.) were added to a solution of alcohol (2.0 mmol) in anhydrous dimethylformamid (DMF) (3 mL). The mixture was stirred at 40 °C for 15 h until the alcohol was completely activated, as observed by the thin-layer chromatography control. The mixture was cooled to room temperature and Boc-L-lysine-OtBu (Novabiochem, 2 mmol) was added at a rate that kept the reaction temperature below 30 °C. Then, the mixture was stirred overnight at room temperature. After the carbamate was formed (TLC control), H_2_O (10 mL) and EtOAc (10 mL) were added to the mixture. The organic layer was separated, and the aqueous layer was extracted with EtOAc (5 mL). The organic layer was subsequently washed with 1 N HCl, H_2_O, and brine, then dried on MgSO_4_. To protect the ncAAs as an oil, the organic solution was concentrated and column chromatography (n-hexane: EtOAC = 1:1) was performed. Then, a 1:1 mixture of TFA:CH_2_Cl_2_ was used for deprotection. After completing deprotection (confirmed by TLC), all volatiles were removed *in vacuo* and any residue was dissolved in methanol. Cold diethyl ether was added to precipitate the pure ncAA, which was filtered out and dried *in vacuo*.

#### S2.1.1 Synthesis of ncAA 2

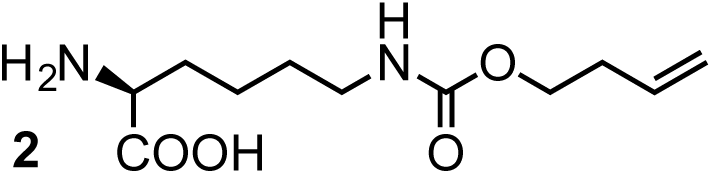

Boc-ncAA-OtBu **2**

^1^H-NMR (CDCl_3_, 400 MHz): δ = 1.23-1.27 (m, 2H), 1.30 (s, 9H), 1.32 (s, 9H), 1.35-1.63 (m, 4H), 2.19-2.24 (m, 2H), 2.99-3.04 (m, 2H), 3.93-4.00 (m, 3H), 4.90-4.98 (m, 2H), 5.16 (br, 2H), 5.59-5.69 (m, 1H).

^13^C-NMR (CDCl_3_, 100 MHz): δ = 171.86, 156.62, 155.39, 134.14, 116.82, 81.50, 79.31, 63.56, 53.70, 46.72, 33.38, 27.83, 22.24.

ncAA **2**

^1^H-NMR (D_2_O, 400 MHz): δ = 1.42-158 (m, 4H), 1.89 (m, 2H), 2.34-2.39 (dd, *J* = 8.0, 4.0 Hz, 1H), 3.13 (t, *J* = 6.0 Hz), 3.89 (t, *J* = 7.5 Hz), 4.06 (m, 2H), 5.07 (m, 2H), 5.84 (m, 1H).

^13^C-NMR (D_2_O, 100 MHz): δ = 174.25, 158.80, 134.80, 117.26, 64.35, 54.45, 39.88, 29.98, 21.56.

HRMS: *m*/*z* calcd for C_20_H_36_N_2_O_6_ [M + Na]+: 400.2573; found:

#### S2.1.2 Synthesis of ncAA 3

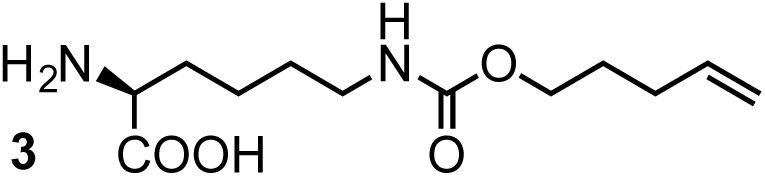

Boc-ncAA-OtBu **3**

^1^H-NMR (CDCl_3_, 400 MHz): δ = 1.30 (s, 9H), 1.32 (s, 9H), 1.34-1.63 (m, 8H), 1.96 (d, 2H, *J* = 4.0 Hz), 3,0 (br, 2H), 4.82 (d, *J* = 8.0 Hz), 4.87 (d, *J* = 16.0 Hz), 5.16 (br, 2H), 5.19 (m, 2H), 5.59-5.69 (m, 1H).

^13^C-NMR (CDCl_3_, 100 MHz): δ = 171.83, 156.71, 155.37, 137.44, 114.93, 81.42, 79.22, 63.84, 53.68, 40.36, 29.85, 28.17, 27.81, 22.24.

ncAA **3**

^1^H-NMR (D_2_O, 400 MHz): δ = 0.99-1.21 (m, 4H), 1.22-1.25 (m, 2H), 1.43 (m, 2H), 1.53 (q, *J* = 16.0 Hz, 1H), 3.59 (m, 3H), 4.55 (d, *J* = 8.0 Hz), 4.60 (d, *J* = 16.0 Hz), 5.37-5.42 (m, 1H).

^13^C-NMR (D_2_O, 100 MHz): ^13^C-NMR (CDCl_3_, 100 MHz): δ = 171.46, 158.41, 137.99, 114.45, 64.54, 52.39, 39.62, 29.07, 28.17, 27.14, 21.18.

#### S2.1.3 Synthesis of ncAA 4

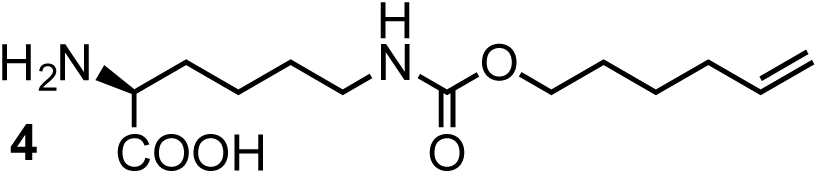

Boc-ncAA-OtBu **4**

^1^H-NMR (CDCl_3_, 400 MHz): δ = 1.21 (s, 9H), 1.24 (s, 9H), 1.32-1.40 (m, 10H), 1.85 (dd, 2H, *J* = 10.0 Hz), 2.91 (br, 2H), 3.81 (t, 2H, *J* = 4.0 Hz), 3.90 (m, 1H), 4.73 (d, *J* = 12.0 Hz), 4.77 (d, *J* = 20.0 Hz), 5.23-5.35 (m, 2H), 5.55 (m, 1H).

^13^C-NMR (CDCl_3_, 100 MHz): δ = 171.79, 156.74, 155.33, 138.07, 114.53, 81.22, 79.02, 64.16, 53.69, 40.23, 33.09, 32.01, 29.24, 28.31, 28.10, 27.72, 24.90, 22.21.

ncAA **4**

^1^H-NMR (D_2_O, 400 MHz): δ = 0.63-0.83 (m, 8H), 1.14-1.25 (m, 4H), 2.32 (m, 2H), 3.23 (m, 3H), 4.13 (d, 2H, *J* = 12.0 Hz), 4.77 (d, 2H, *J* = 20.0 Hz), 5.02 (m, 1H).

^13^C-NMR (D_2_O, 100 MHz): δ = 170.75, 157.88, 137.79, 113.32, 64.53, 52.02, 30.02,
27.02, 23.87, 20.93.

#### S2.1.4 Synthesis of ncAA 5

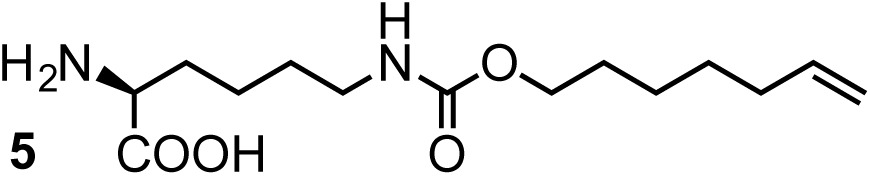

Boc-ncAA-OtBu **5**

^1^H-NMR (CDCl_3_, 400 MHz): δ = 1.31 (s, 9H), 1.33 (s, 9H), 1.41-1.49 (m, 12H), 1.92 (dd, 2H, *J* = 6.0 Hz), 3.02 (br, 2H), 3.91(t, 2H, *J* = 6.0 Hz), 4.01 (br, 1H), 4.80 (d, *J* = 9.0 Hz), 4.86 (d, *J* = 16.0 Hz), 5.16 (br, 2H), 5.67 (m, 1H).

^13^C-NMR (CDCl_3_, 100 MHz): δ = 171.86, 156.80, 155.38, 138.50, 114.34, 81.41, 79.28, 64.53, 53.68, 40.40, 33.48, 32.27, 29.35, 28.78, 28.38, 28.20, 27.84, 25.21, 22.25.

ncAA **5**

^1^H-NMR (D_2_O, 400 MHz): δ = 0.54-0.80 (m, 12H), 1.22 (m, 2H), 2.32 (m, 2H), 3.26 (m, 3H), 4.12 (d, 2H, *J* = 20.0 Hz), 4.20 (d, 2H, *J* = 16.0 Hz), 4.63 (m, 1H).

^13^C-NMR (D_2_O,100 MHz): δ = 171.05, 158.05, 138.67, 113.12, 64.79, 52.13, 39.31, 32.27,28.77, 27.77, 27.20, 23.98, 20.85.

#### S2.1.5 Synthesis of ncAA 6

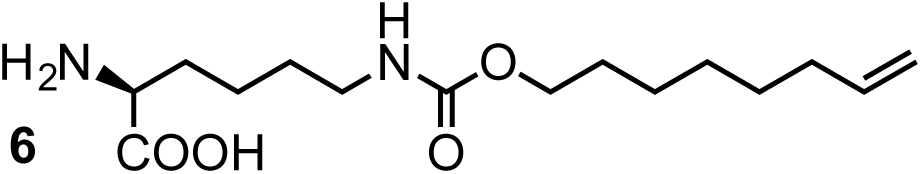

Boc-ncAA-OtBu **6**

^1^H-NMR (CDCl_3_, 400 MHz): δ = 1.21 (s, 9H), 1.23 (s, 9H), 1.15-1.37 (m, 14H), 1.81 (dd, 2H, *J* = 6.0 Hz), 2.92 (br, 2H), 3.80 (M, 2H), 3.90 (m, 1H), 4.70 (d, *J* = 12.0 Hz), 4.75 (d, *J* = 16.0 Hz), 5.24-5.35 (m, 2H), 5.52-5.58 (m, 1H).

^13^C-NMR (CDCl_3_, 100 MHz): δ = 171.79, 156.76, 155.33, 138.49, 114.12, 81.17, 78.98, 64.33, 53.68, 40.21, 33.42, 32.01, 29.24, 28.72, 28.52, 27.71, 25.60, 22.20.

ncAA **6**

^1^H-NMR (D_2_O, 400 MHz): δ = 0.26-0.68 (m, 14H), 1.15 (m, 2H), 2.24 (m, 2H), 3.15 (m, 3H), 4.03 (d, 2H, *J* = 12.0 Hz), 4.10 (d, 2H, *J* = 16.0 Hz), 4.91-4.97 (m, 1H).

^13^C-NMR (D_2_O, 100 MHz): δ = 170.86, 158.04, 138.63, 113.12, 65.13, 52.47, 39.11, 32.09, 28.77, 27.70, 27.30, 24.60, 20.83.

#### S2.1.6 Synthesis of ncAA 7

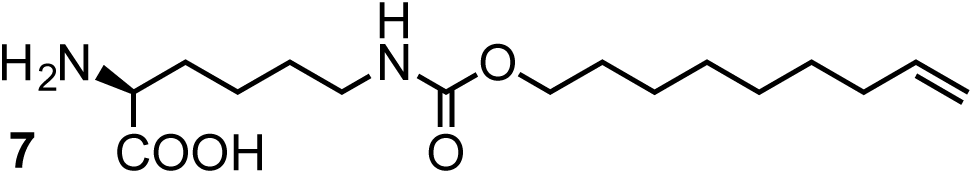

Boc-ncAA-OtBu **7**

^1^H-NMR (CDCl_3_, 400 MHz): δ = 1.18 (s, 9H), 1.24 (s, 9H), 1.12-1.37 (m, 16H), 1.83 (dd, 2H, *J* = 6.0 Hz), 2.96 (m, 2H), 3.82 (m, 2H), 3.93 (m, 1H), 4.72 (d, *J* = 12.0 Hz), 4.78 (d, *J* = 16.0 Hz), 5.25-5.30 (m, 2H), 5.55-5.60 (m, 1H).

^13^C-NMR (CDCl_3_, 100 MHz): δ = 171.80, 156.77, 155.34, 138.63, 114.07, 81.23, 79.03, 64.40, 53.68, 40.25, 33.51, 32.54, 29.26, 28.92, 28.76, 28.58, 28.12, 27.74, 25.61, 22.21.

ncAA **7**

^1^H-NMR (D2O, 400 MHz): δ = 0.60-0.88 (m, 14H), 1.21 (m, 4H), 2.40 (m, 2H), 3.29 (m, 3H), 3.34 (t, 1H, *J* = 8.0 Hz), 4.19 (d, 2H, *J* = 8.0 Hz), 4.26 (d, 2H, *J* = 20.0 Hz), 5.04-5.10 (m, 1H).

^13^C-NMR (D2O, 100 MHz): δ = 171.05, 158.04, 138.19, 113.11, 64.80, 52.21, 39.46, 32.83, 28.83, 28.24, 28.05, 28.00, 27.94, 27.86, 24.84, 21.07.

#### S2.1.7 Synthesis of ncAA 8

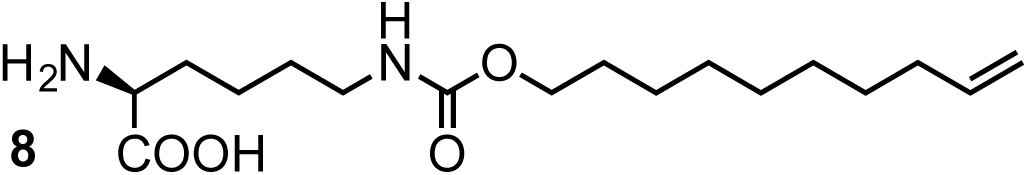

Boc-ncAA-OtBu **8**

^1^H-NMR (CDCl_3_, 400 MHz): δ = 1.29 (s, 9H), 1.34 (s, 9H), 1.15-1.37 (m, 18H), 1.88 (dd, 2H, *J* = 6.0 Hz), 3.00 (m, 2H), 3.87 (m, 2H), 3.97 (m, 1H), 4.76 (d, *J* = 8.0 Hz), 4.82 (d, *J* = 16.0 Hz), 5.22 (m, 2H), 5.58-5.68 (m, 1H).

^13^C-NMR (CDCl_3_, 100 MHz): δ = 171.85, 156.84, 155.39, 138.77, 114.05, 81.38, 79.20, 64.55, 62.31, 53.68, 40.32, 33.59, 29.31, 29.29, 29.20, 29.08, 28.91, 28.85, 28.70, 28.16, 27.79, 25.69, 22.23.

ncAA **8**

^1^H-NMR (D2O, 400 MHz): δ = 0.7-1.06 (m, 16H), 1.42-1.49 (m, 4H), 2.56 (m, 2H), 3.45 (m, 2H), 3.51 (t, 1H, *J* = 8.0 Hz), 4.35 (d, 2H, *J* = 8.0 Hz), 4.41 (d, 2H, *J* = 16.0 Hz), 5.17-5.27 (m, 1H).

^13^C-NMR (D2O, 100 MHz): δ = 171.24, 158.04, 138.12, 113.37, 65.02, 52.23, 39.86, 33.15, 28.79, 28.66, 28.49, 28.31, 28.20, 25.01, 21.26.

#### S2.1.8 Synthesis of ncAA 9

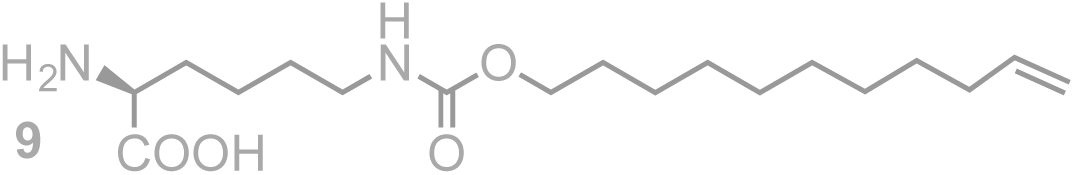

Boc-ncAA-OtBu **9**

^1^H-NMR (CDCl_3_, 400 MHz): δ = 1.21 (s, 9H), 1.28 (s, 9H), 1.52 (m, 20H), 1.96 (dd, 2H, *J* = 6.0 Hz), 3.08 (m, 2H), 3.95 (m, 2H), 4.07 (m, 1H), 4.85 (d, *J* = 8.0 Hz), 4.86 (d, *J* = 16.0 Hz), 5.15 (m, 1H), 5.76-5.78 (m, 1H).

^13^C-NMR (CDCl_3_, 100 MHz): δ = 171.89, 156.83, 155.41, 138.99, 114.07, 81.61, 79.42, 64.72, 53.68, 40.47, 33.59, 29.60, 29.39, 29.31, 29.19, 29.00, 28.81, 28.25, 28.70, 27.90, 27.79, 25.69, 21.95.

ncAA **9**

^1^H-NMR (D2O, 400 MHz): δ = −0.18-0.4 (m, 18H), 0.68 (m, 4H), 2.56 (m, 2H), 1.82 (m, 2H), 2.71 (m, 3H), 3.57 (d, 2H, *J* = 12.0 Hz), 3.65 (d, 2H, *J* = 16.0 Hz), 5.17-5.27 (m, 1H).

^13^C-NMR (D2O, 100 MHz): δ = 170.34, 157.91, 138.93, 111.94, 64.99, 51.84, 38.79, 31.94, 28.17, 27.42, 27.35, 27.22, 27.12, 27.01, 26.84, 23.87, 20.34.

#### S2.1.9 Synthesis of ncAA 10

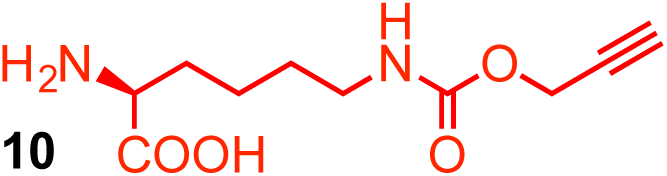

Boc-ncAA-OtBu **10**

^1^H-NMR (CDCl_3_, 400 MHz): δ = 1.31 (s, 9H), 1.33 (s, 9H), 1.57 (m, 6H), 2.47 (m, 2H), 3.20 (m, 2H), 4.15 (m, 1H), 4.68 (br, 2H), 4.97 (br, 1H), 5.11 (br, 1H).

^13^C-NMR (CDCl_3_, 100 MHz): δ = 171.87, 156.18, 155.41, 81.67, 80.34, 79.49, 69.55, 62.36 53.67, 40.55, 33.59, 29.60, 28.27, 27.92, 27.81, 22.27, 19.32.

ncAA **10**

^1^H-NMR (D_2_O, 400 MHz): δ = 1.31 (m, 2H), 1.45 (m, 2H), 1.78 (m, 2H), 2.78 (m, 1H), 3.05 (t, 2H, *J* = 6.0 Hz), 3.67(t, 2H, *J* = 6.0 Hz), 4.56 (s, 2H).

^13^C-NMR (D_2_O, 100 MHz): δ = 174.34, 157.68, 78.81, 75.53, 54.72, 52.25, 40.09, 30.01, 28.16, 21.53.

#### S2.1.10 Synthesis of ncAA 11

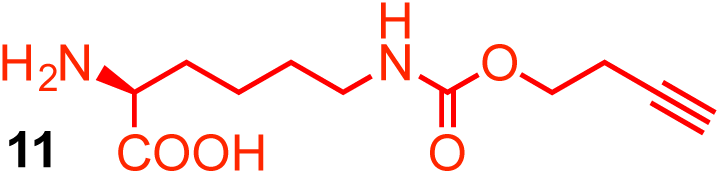

Boc-ncAA-OtBu **11**

^1^H-NMR (CDCl_3_, 400 MHz): δ = 1.31 (s, 9H), 1.33 (s, 9H), 1.27-1.53 (m, 6H), 1.94 (t, 1H, *J* = 8.0 Hz), 2.43 (t, 2H, *J* = 6.0 Hz), 3.08 (m, 2H), 4.07 (m, 3H), 5.09 (br, 2H).

^13^C-NMR (CDCl_3_, 100 MHz): δ = 171.87, 156.18, 155.41, 81.67, 80.34, 79.49, 69.55, 62.36 53.67, 40.55, 33.59, 29.60, 28.27, 27.92, 27.81, 22.27, 19.32.

ncAA**11**

^1^H-NMR (D_2_O, 400 MHz): δ = 1.26-1.39 (m, 4H), 1.68-1.75 (m, 2H), 2.28 (m, 1H), 2.43 (m, 2H), 3.02 (m, 2H), 3.63 (m, 1H), 4.03 (m, 2H).

^13^C-NMR (D_2_O, 100 MHz): δ = 174.59, 158.36, 82.02, 70.79, 62.68, 54.60, 40.14, 30.04, 28.56, 21.55, 18.66.

#### S2.1.11 Synthesis of ncAA 12

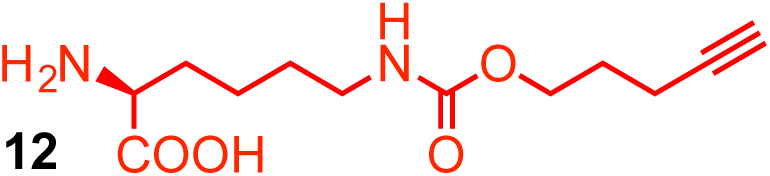

Boc-ncAA-OtBu **12**

^1^H-NMR (CDCl_3_, 400 MHz): δ = 1.31 (s, 9H), 1.33 (s, 9H), 1.41-1.59 (m, 6H), 1.56-1.59 (m, 2H), 1.68 (m, 1H), 2.21 (t, 2H, *J* = 6.0 Hz), 3.10 (t, 2H, *J* = 6.0 Hz), 4.09 (m, 3H), 4.92-5.09 (br, 2H).

^13^C-NMR (CDCl_3_, 100 MHz): δ = 171.87, 156.53, 155.43, 83.15, 81.73, 79.54, 68.89, 63.20, 53.66, 40.63, 32.53, 28.28, 27.94, 22.30, 15.10.

ncAA **12**

^1^H-NMR (D_2_O, 400 MHz): δ = 1.28-1.41 (m, 4H), 1.70-1.74 (m, 4H), 2.22 (m, 2H), 2.23 (m, 1H), 3.01 (m, 2H), 3.63 (t, 2H, *J* = 6.0 Hz), 4.04 (t, 2H, *J* = 6.0 Hz).

^13^C-NMR (D_2_O, 100 MHz): δ = 174.25, 158.77, 84.88, 69.55, 64.09, 54.50, 39.87, 30.00, 28.52, 27.21, 21.56, 14.32.

#### S2.1.12 Synthesis of ncAA 13

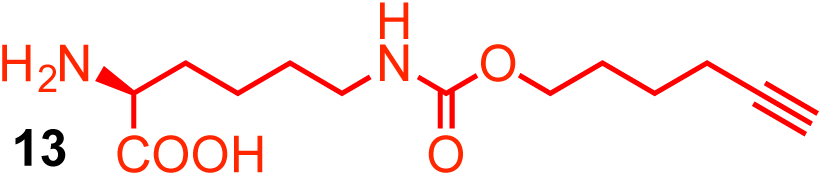

Boc-ncAA-OtBu **13**

^1^H-NMR (CDCl_3_, 400 MHz): δ = 1.19 (s, 9H), 1.21 (s, 9H), 1.26-1.35 (m, 10H), 1.78 (m, 1H), 1.96 (m, 2H), 2.90 (m, 2H), 3.81(m, 2H), 3.87 (m, 1H), 5.23 (m, 1H), 5.39 (br, 1H).

^13^C-NMR (CDCl_3_, 100 MHz): δ = 171.77, 156.63, 155.31, 83.58, 81.18, 78.99, 68.81, 63.68, 53.68, 40.22, 28.09, 27.90, 27.71, 24.64, 22.21, 17.79.

ncAA **13**

^1^H-NMR (D_2_O, 400 MHz): δ = 0.98-1.24 (m, 8H), 1.41 (m, 2H), 1.51 (m, 2H), 1.75 (t, 2H, *J* = 4.0 Hz), 1.77 (m, 1H), 2.66 (t, 2H, *J* = 6.0 Hz), 3.60 (m, 2H).

^13^C-NMR (D_2_O, 100 MHz): δ = 171.45, 158.44, 85.55, 68.87, 64.68, 52.38, 39.59, 28.10, 27.05, 23.93, 21.15, 16.88.

#### S2.1.13 Synthesis of ncAA 14

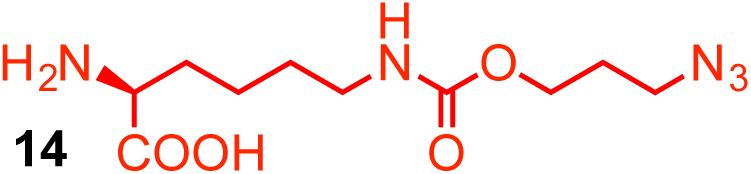

Boc-ncAA-OtBu **14**

^1^H-NMR (CDCl_3_, 400 MHz): δ = 1.29 (s, 9H), 1.32 (s, 9H), 1.18-1.47 (m, 6H), 1.64 (m, 1H), 3.03 (dd, 2H, *J* = 8.0 Hz), 4.01 (m, 3H), 5.18 (m, 2H).

^13^C-NMR (CDCl_3_, 100 MHz): δ = 171.81, 156.38, 155.41, 81.53, 79.33, 61.39, 60.20, 53.68, 48.10, 40.47, 28.47, 28.18, 27.82, 22.27, 20.83.

ncAA **14**

^1^H-NMR (D_2_O, 400 MHz): δ = 1.42-1.60 (m, 4H), 1.94 (m, 4H), 3.17 (m, 2H), 3.46 (m, 2H), 3.93 (m, 1H), 4.17 (m, 2H).

^13^C-NMR (D_2_O, 100 MHz): δ = 173.60, 159.12, 65.98, 62.64, 53.88, 48.02, 39.89, 27.65, 27.01, 21.50.

#### S2.1.14 Synthesis of ncAA 18

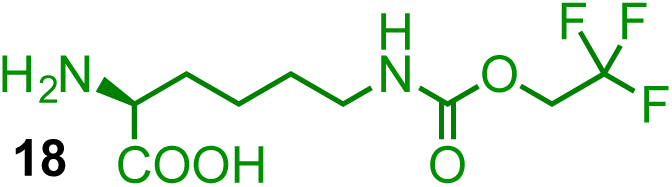

Boc-ncAA-OtBu **18**

^1^H-NMR (CDCl_3_, 400 MHz): δ = 1.28 (s, 9H), 1.30 (s, 9H), 1.38 (m, 6H), 1.78 (m, 1H), 3.04 (dd, 2H, *J* = 6.0 Hz), 3.98 (m, 1H), 4.30 (dd, 2H, *J* = 8.0 Hz), 5.24 (m, 1H), 5.89 (m, 1H).

^13^C-NMR (CDCl_3_, 100 MHz): δ = 171.84, 155.47, 154.48, 121.72, 124.47, 81.49, 79.30, 60.24, 60.15, 59.88, 53.65, 40.66, 28.89, 28.01, 27.65, 27.39, 22.12.

ncAA **18**

^1^H-NMR (D_2_O, 400 MHz): δ = 0.65-0.77 (m, 4H), 1.11-1.20 (m, 2H), 2.36 (t, 2H, *J* = 4.0 Hz), 3.25 (t, 2H, *J* = 6.0 Hz), 3.71 (q, 2H, *J* = 8.0 Hz).

^13^C-NMR (D_2_O, 100 MHz): δ = 170.87, 155.70, 52.08, 39.40, 28.65, 27.47, 20.51.

#### S2.1.15 Synthesis of ncAA 19

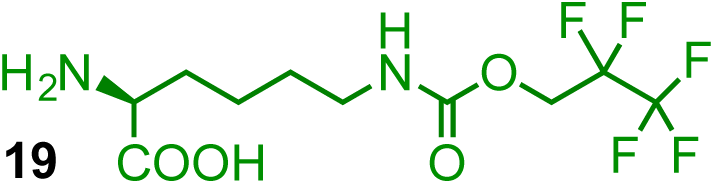

Boc-ncAA-OtBu **19**

^1^H-NMR (CDCl_3_, 400 MHz): δ = 1.31 (s, 9H), 1.33 (s, 9H), 1.38-1.65 (m, 8H), 3.08 (dd, 2H, *J* = 6.0 Hz), 4.03 (m, 1H), 4.45 (dd, 2H, *J* = 12.0 Hz), 5.20 (m, 1H), 5.76 (m, 1H).

^13^C-NMR (CDCl_3_, 100 MHz): δ = 171.86, 155.48, 154.53, 81.57, 79.37, 60.24, 60.15, 59.88, 53.63, 40.76, 28.89, 28.01, 27.62, 27.39, 22.13.

ncAA **19**

^1^H-NMR (D_2_O, 400 MHz): δ = 0.65-0.77 (m, 4H), 1.11-1.20 (m, 2H), 2.36 (t, 2H, *J* = 4.0 Hz), 3.25 (t, 2H, *J* = 6.0 Hz), 3.71 (q, 2H, *J* = 8.0 Hz).

^13^C-NMR (D_2_O, 100 MHz): δ = 170.87, 155.70, 52.08, 39.40, 28.65, 27.47, 20.51.

#### S2.1.16 Synthesis of ncAA 20

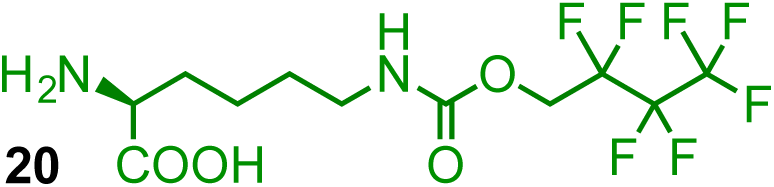

Boc-ncAA-OtBu **20**

^1^H-NMR (CDCl_3_, 400 MHz): δ = 1.31 (s, 9H), 1.33 (s, 9H), 1.38-1.65 (m, 8H), 3.08 (dd, 2H, *J* = 6.0 Hz), 4.03 (m, 1H), 4.45 (dd, 2H, *J* = 12.0 Hz), 5.20 (m, 1H), 5.76 (m, 1H).

^13^C-NMR (CDCl_3_, 100 MHz): δ = 171.86, 155.48, 154.53, 81.57, 79.37, 60.24, 60.15, 59.88, 53.63, 40.76, 28.89, 28.01, 27.62, 27.39, 22.13.

^1^H-NMR (D_2_O, 400 MHz): δ = 0.81-0.94 (m, 4H), 1.33 (m, 2H), 2.52 (t, 2H, *J* = 8.0 Hz), 3.42 (m, 2H), 3.71 (t, 2H, *J* = 12.0 Hz).

^13^C-NMR (D_2_O, 100 MHz): δ = 171.47, 155.92, 51.77, 39.43, 28.89, 27.12, 20.76.

#### S2.1.17 Synthesis of ncAA 21

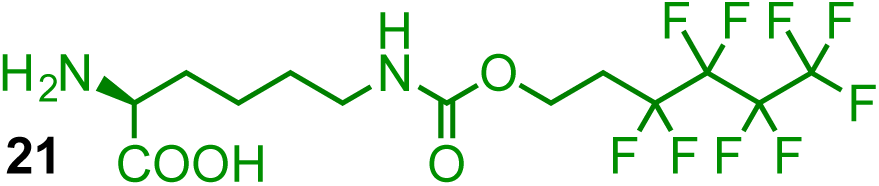

Boc-ncAA-OtBu **21**

^1^H-NMR (CDCl_3_, 400 MHz): δ = 1.37 (s, 9H), 1.39 (s, 9H), 1.31-1.56 (m, 6H), 2.27 (M, 2H), 3.11 (m, 2H), 4.03 (m, 1H), 4.30 (m, 2H), 5.17 (m, 1H), 5.24 (m, 1H).

^13^C-NMR (CDCl_3_, 100 MHz): δ = 171.87, 155.88, 154.53, 81.67, 79.48, 56.42, 53.64, 40.56, 28.89, 28.01, 27.74, 22.23.

ncAA **21**

^1^H-NMR (D_2_O, 400 MHz): δ = 0.60-0.73 (m, 4H), 1.11 (m, 2H), 1.63 (m, 2H), 2.29 (t, 2H, *J* = 6.0 Hz), 3.20 (m, 1H), 3.50 (t, 2H, *J* = 8.0 Hz).

^13^C-NMR (D_2_O, 100 MHz): δ = 171.46, 157.06, 56.67, 51.77, 39.46, 28.67, 27.10, 20.81.

#### S2.1.18 Synthesis of ncAA 22

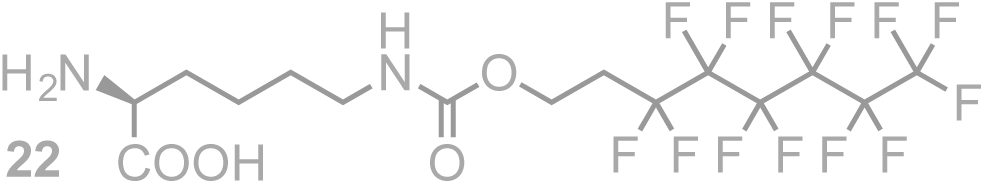

Boc-ncAA-OtBu **22**

^1^H-NMR (CDCl_3_, 400 MHz): δ = 1.26 (s, 9H), 1.28 (s, 9H), 1.38 (m, 6H), 2.24-2.35 (m, 2H), 3.00 (m, 2H), 3.98 (m, 1H), 4.18 (m, 2H), 5.28 (m, 1H), 5.58 (m, 1H).

^13^C-NMR (CDCl_3_, 100 MHz): δ = 171.88, 155.91, 155.50, 81.66, 79.48, 56.44, 53.64, 40.55, 32.41, 30.96, 30.74, 30.53, 29.12, 28.04, 27.70, 22.23.

ncAA **22**

^1^H-NMR (D_2_O, 400 MHz): δ = 0.36-0.45 (m, 4H), 0.85-0.92 (m, 2H), 1.36 (m, 2H), 2.05 (m, 2H), 2.96 (m, 1H), 3.26 (m, 2H).

^13^C-NMR (D_2_O, 100 MHz): δ = 170.69, 156.87, 56.33, 51.78, 39.08, 28.41, 27.11, 20.46.

#### S2.1.19 Synthesis of ncAA 23

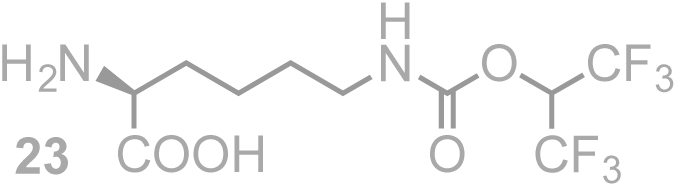

Boc-ncAA-OtBu **23**

^1^H-NMR (CDCl_3_, 400 MHz): δ = 1.27 (s, 9H), 1.29 (s, 9H), 1.35-1.60 (m, 6H), 1.97 (dd, 2H, *J* = 4.0 Hz), 3.01 (m, 3H), 4.03 (m, 2H), 5.21 (m, 1H), 5.47 (m, 1H).

^13^C-NMR (CDCl_3_, 100 MHz): δ = 171.80, 155.95, 155.44, 81.41, 79.22, 60.50, 53.67, 40.40, 32.12, 27.95, 27.58, 22.19.

ncAA **23**

^1^H-NMR (D_2_O, 400 MHz): δ = 1.35-1.48 (m, 4H), 1.78 (m, 2H), 2.09 (m, 2H), 3.03 (t, 2H, *J* = 6.0 Hz), 3.95 03 (t, 2H, *J* = 8.0 Hz), 4.10 (m, 2H).

^13^C-NMR (D_2_O, 100 MHz): δ = 172.17, 158.22, 66.97, 52.82, 39.78, 29.35, 28.31, 22.81, 21.38, 16.91.

### S2.2 Synthesis of ncAAs with Acid-Sensitive Carbamates (ncAAs 28 and 29)

#### S2.2.1 General Procedure

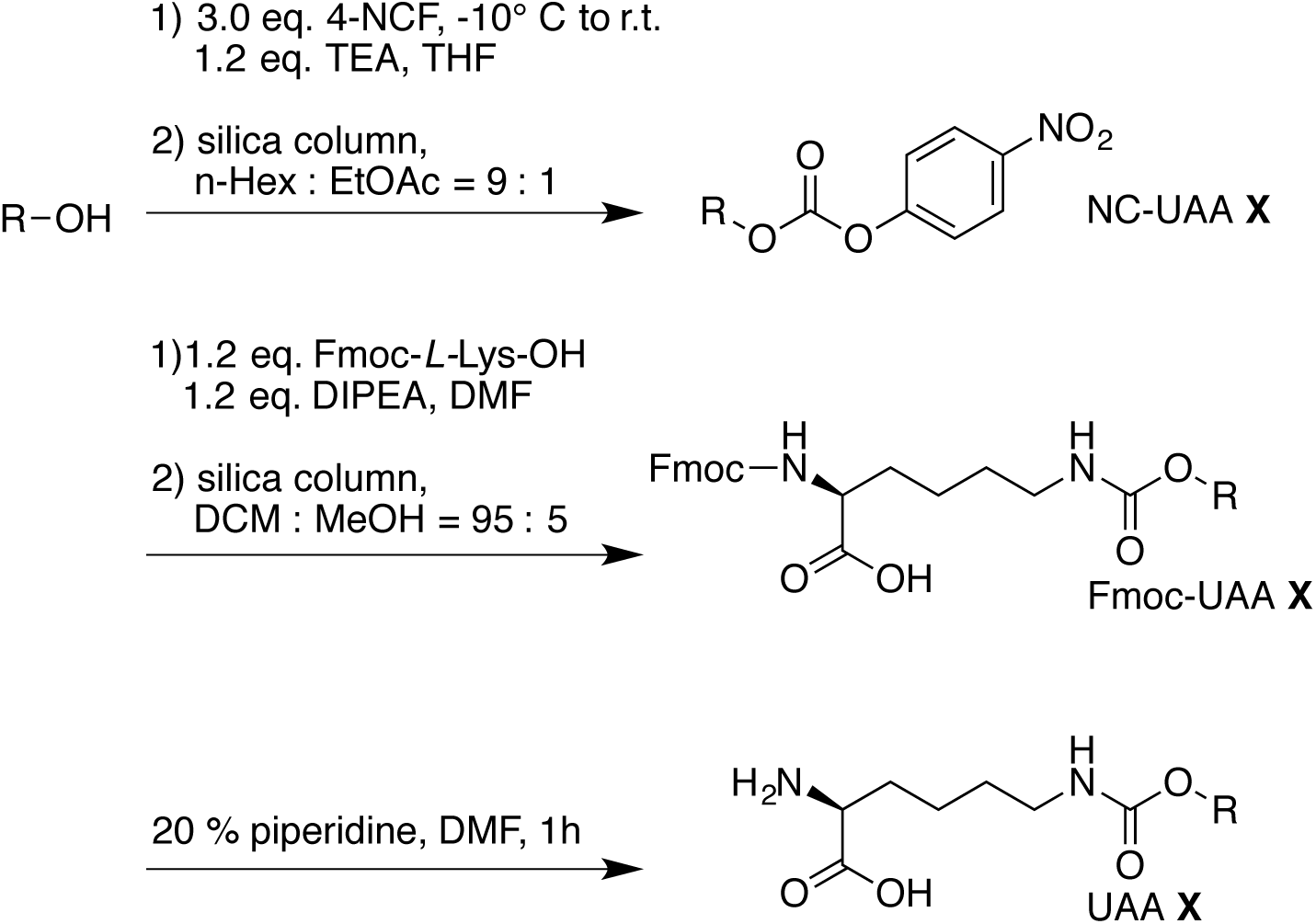

#### S2.2.2 Synthesis of the Nitrophenol-Carbamate Precursor (NC-ncAA)

Alcohol (1.54 g, 10.0 mmol) and TEA (1.70 ml, 12 mmol, 1.2 eq) were dissolved in THF (40 ml) and dripped into a stirred solution of 4-nitrophenyl chloroformate (4-NCF, 6.04 g, 30 mmol, 3.0 eq) in THF (36 ml) over a period of 1 h at −10 °C. The reaction mixture was allowed to warm to room temperature and was stirred overnight. THF was removed *in vacuo,* and water was added to the residue. The mixture was extracted three times with EtOAc. The collected organic phase were dried on MgSO_4_ following by column chromatography (n-hexane : EtOAc = 9 : 1) to yield the desired yellow-oil product.

#### S2.2.3 Synthesis of the Fmoc-Protected ncAA (Fmoc-ncAA)

Fmoc-L-Lys-OH (Novabiochem, 0.69 g, 1.87 mmol, 1.2 eq.) was suspended under argon in anhydrous DMF (0.2 M, 8 ml) containing DIEA (0.24 g, 0.33 ml, 1.87 mmol, 1.2 eq.). To this white suspension, a clear solution of the nitrophenol-carbamate precursor (1.56 mmol, 1.0 eq.) in anhydrous DMF (0.2 M, 8 ml) was added dropwise under argon at room temperature over a period of 2 h. The reaction mixture was stirred for an additional 4 h at room temperature. H_2_O (50 ml) and EtOAc (150 ml) were added and the aqueous layer was adjusted to a pH range of 1–3 with 1 N HCl. The phases were separated and the aqueous layer was extracted with EtOAc (2 x 50 ml). The organic layers were washed with saturated NaCI solution (2 x 50 ml) and dried on Na_2_SO_4_. All volatiles were evaporated under reduced pressure and the crude product was purified by column chromatography (DCM : MeOH 95 : 5 v/v) to yield the Fmoc-protected ncAA as a white solid.

#### S2.2.4 Deprotection of the Fmoc-ncAA

The Fmoc-protected ncAA was dissolved in 20% piperidine in DMF (20 ml per mmol of ncAA) and stirred for 1 h at room temperature. All volatiles were removed under reduced pressure to yield the pure compound as a solid.

### S2.3.1 Synthesis of ncAA 28

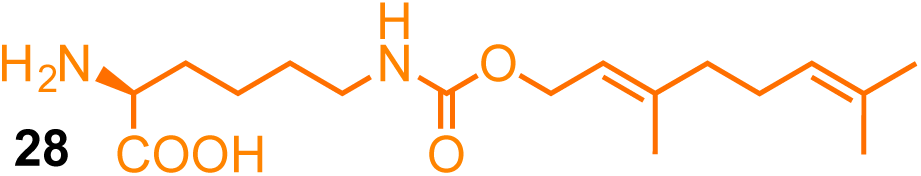

NC-ncAA **28**

^1^H-NMR (CDCl_3_, 400 MHz): δ = 1.63 (s, 3H), 1.70 (s, 3H), 1.83 (s, 3H), 2.15 (m, 4H), 4.78 (d, 2H, *J* = 4.0 Hz), 5.10 (m, 1H), 5.47 (t, 1H, *J* = 8.0 Hz), 7.40 (d, 2H, *J* = 12.0 Hz), 8.29 (d, 2H, *J* = 8.0 Hz).

Fmoc-ncAA **28**

^1^H-NMR (CDCl_3_, 400 MHz): δ = 1.62 (s, 3H), 1.70 (s, 3H), 1.76 (s, 3H), 1.44-1.91 (m, 6H), 2.09 (m, 2H), 3.16 (m, 2H), 4.41 (m, 2H), 4.59 (m, 2H), 5.11 (m, 2H), 5.35 (m, 1H), 5.97 (d, *J* = 4.0 Hz), 7.31 (t, 2H, *J* = 8.0 Hz), 7.39 (t, 2H, *J* = 6.0 Hz), 7.55-7.63 (m, 2H), 7.66 (d, *J* = 8.0 Hz).

^13^C-NMR (CDCl_3_, 100 MHz): δ = 176.25, 176.03, 175.63, 171.51, 157.13, 156.44, 143.96, 143.78, 142.14, 141.34, 132.12, 127.13, 123.69, 120.00, 67.14, 60.57, 53.70, 47.21, 40.63, 32.20, 29.41, 26.72, 25.72, 22.38, 21.07, 20.79, 17.70, 14.22.

ncAA **28**

^1^H-NMR (D_2_O, 400 MHz): δ = 1.64 (s, 3H), 1.71 (s, 3H), 1.75 (s, 3H), 1.20-1.91 (m, 6H), 2.16 (m, 2H), 3.15 (m, 2H), 3.75 (m, 1H), 4.55 (m, 2H), 4.59 (m, 2H), 5.42 (m, 1H).

#### S2.3.2 Synthesis of ncAA 29

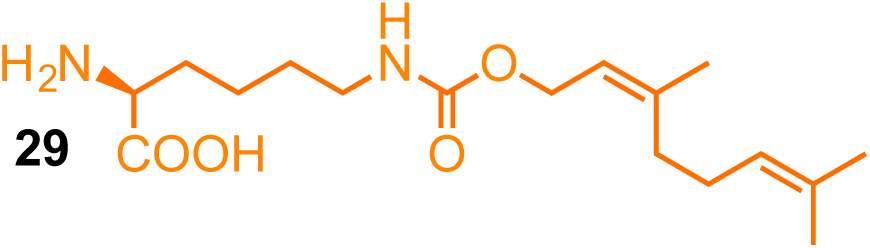

NC-ncAA **29**

^1^H-NMR (CDCl_3_, 400 MHz): δ = 1.62 (s, 3H), 1.70 (s, 3H), 1.78 (s, 3H), 2.12 (m, 4H), 4.82 (d, 2H, *J* = 8.0 Hz), 5.10 (m, 1H), 5.47 (t, 1H, *J* = 8.0 Hz), 7.40 (d, 2H, *J* = 8.0 Hz), 8.29 (d, 2H, *J* = 8.0 Hz).

Fmoc-ncAA **29**

^1^H-NMR (CDCl_3_, 400 MHz): δ = 1.61 (s, 3H), 1.64 (s, 3H), 1.69 (s, 3H), 1.43-1.91 (m, 6H), 2.07 (m, 2H), 3.20 (m, 2H), 4.22 (m, 1H), 4.40 (m, 2H), 4.66-4.82 (m, 2H), 5.09 (m, 2H), 5.33 (m, 1H), 7.29 (t, 2H, *J* = 6.0 Hz), 7.38 (t, 2H, *J* = 6.0 Hz), 7.61 (m, 2H), 7.76 (d, *J* = 8.0 Hz).

^13^C-NMR (CDCl_3_, 100 MHz): δ = 176.03, 175.20, 157.13, 156.35, 144.01, 143.88, 142.58, 141.29, 131.79, 127.07, 123.76, 119.96, 67.23, 62.00, 53.88, 47.15, 39.54, 31.73, 26.31, 25.69, 22.56, 17.70, 16.47.

ncAA **29**

^1^H-NMR (D_2_O, 400 MHz): δ = 1.60 (s, 3H), 1.66 (s, 3H), 1.70 (s, 3H), 1.46-1.89 (m, 6H), 2.05 (m, 2H), 3.11 (m, 2H), 3.67 (m, 1H), 4.55 (m, 2H), 5.32 (m, 1H).

